# SHP2 binds directly to SOS1 to enable RAS activation

**DOI:** 10.64898/2026.06.12.731952

**Authors:** Toshiyuki Araki, Yashika Agrawal, Sachin S. Katti, Hiep L.D. Nguyen, Daniel Pushparaju Yeggoni, Mitchell J. Geer, Wei Wei, Daniëlle T.J. Woutersen, Tieme Bijlsma, Cameron Genxuan Lian, Natalie M. Clark, Namrata D. Udeshi, Steven A. Carr, Heidi Stuhlmann, Michael A. Davies, Eli Rothenberg, Jeroen den Hertog, Rebecca Page, Benjamin G. Neel, Wolfgang Peti

## Abstract

The protein-tyrosine phosphatase SHP2 (*PTPN11*) regulates growth factor- and cytokine-induced RAS/ERK MAP kinase (MAPK) pathway activation, and aberrant SHP2 function causes developmental disorders and cancer^1–5^. It is widely believed that the catalytic activity of SHP2 is essential for pathway activation^1,4,6–8^. This view has shaped our interpretation of how germline *PTPN11* mutations cause Noonan Syndrome (NS) and NS with Multiple Lentigines (NS-ML)^2,9^ and how somatic mutations contribute to myeloproliferative neoplasms and solid tumors^1^. Here we identify a previously undetected, protein-tyrosine phosphatase (PTP) activity-independent mechanism that revises our understanding of how SHP2 promotes RAS/ERK activation. We find that certain mutations of the nucleophilic cysteine that abolish catalytic activity still promote RAS/ERK pathway activation in normal and neoplastic mammalian cells, zebrafish embryos, and mice. Structural studies show that the SHP2 PTP domain binds directly to the Son of Sevenless 1 (SOS1) Dbl homology (DH) domain. Proximity labeling and super-resolution microscopy demonstrate that SHP2/SOS1 interaction occurs in cells and facilitates SOS1 translocation to the plasma membrane to form clusters. Our results overturn decades of dogma on SHP2 regulation of the RAS/ERK pathway and provide new insights into the mechanism of action of disease-associated *PTPN11* mutations.

## Main

The RAS/ERK MAP kinase (RAS/ERK) pathway is activated by most receptor tyrosine kinases (RTKs), including members of the EGF (EGFR), platelet-derived growth factor (PDGFR), and fibroblast growth factor (FGFR) receptor families, as well as by many cytokine receptors and integrins^10–12^. The typical pathway module comprises a receptor coupled to RAS/RAF/MEK/ERK. RAS GTPases (KRAS, NRAS, HRAS) cycle between RAS•GTP “on” and RAS•GDP “off” states, gated by guanine nucleotide exchange factors (RAS-GEFs) and GTPase-activating proteins (RAS-GAPs)^13,14^. RAS-GTP recruits RAF kinases (BRAF/RAF1/ARAF)^15,16^ to the plasma membrane (PM) via RAS-binding (RBD) and cysteine-rich (CRD) domains^17^, where they dimerize and activate MEK1/2, which then activate ERK1/2^12,17,18^. ERK1/2 phosphorylate downstream kinases, transcription factors, and cytoskeletal components to regulate survival, proliferation, differentiation, and migration depending on the cellular context^19^. Although in some circumstances, RAS-GTP can engage other effectors (e.g., PI3K p110, RAL-GDS, TIAM1)^13,20^, in normal cells its main role is to activate the RAF/MEK/ERK cascade^21^. Aberrant RAS/ERK signaling causes “RASopathies”^2,9^, benign tumors (e.g., neurofibromatosis type I, metachondromatosis^22,23^), temporal lobe epilepsy^3^, and cancer^10,12,13,24^. Consequently, RAS/ERK pathway components are major therapeutic targets, and understanding the molecular details of pathway regulation is essential to human health, disease, and therapeutics.

SHP2 *(PTPN11*) is a key regulator of RAS, essential for normal development, a major RASopathy gene, a context-specific proto-oncogene or tumor suppressor, and the target of allosteric inhibitors (SHP2is) in clinical trials^1,25,26^. *Ptpn11^-/-^* mice die peri-implantation owing to trophoblast stem cell death/insufficiency^27^. Germline *PTPN11* mutations cause two phenotypically related RASopathies, NS and NS-ML^2,9,28^. NS mutants typically map to the N-SH2 or PTP domain and shift SHP2 to its “open”, active state^29–33^, thereby increasing PTP activity and RAS/ERK pathway output^30,31,34,35^ (**Fig. 1a**). By contrast, NS-ML alleles alter the PTP domain, and while they too increase open state occupancy, they are catalytically impaired^36–38^. NS-ML pathogenesis was initially thought to reflect increased PI3K/AKT activation^36^, but recent studies show that NS-ML mutants can also increase ERK activation, purportedly by prolonging binding to upstream regulators ^37–39^. Somatic gain-of-function *PTPN11* mutations (along with some strong NS alleles) are the most common cause of juvenile myelomonocytic leukemia (JMML)^40,41^, a potentially fatal childhood myeloproliferative neoplasm (MPN) characterized by cytokine-independent proliferation of myeloid progenitors^42^. Similar mutations contribute to some cases of pediatric B-cell acute lymphoblastic leukemia, neuroblastoma, some adult leukemias, and several solid tumors, especially in the setting of therapy resistance^41,43–47^. Remarkably, recent studies also implicate somatic *Ptpn11* mutations in the pathogenesis of some cases of temporal lobe epilepsy^3^. In addition, wild-type (WT) SHP2 (SHP2^WT^) is required for “adaptive resistance” to RAS/ERK pathway inhibitors^48–55^.

**Fig. 1.**
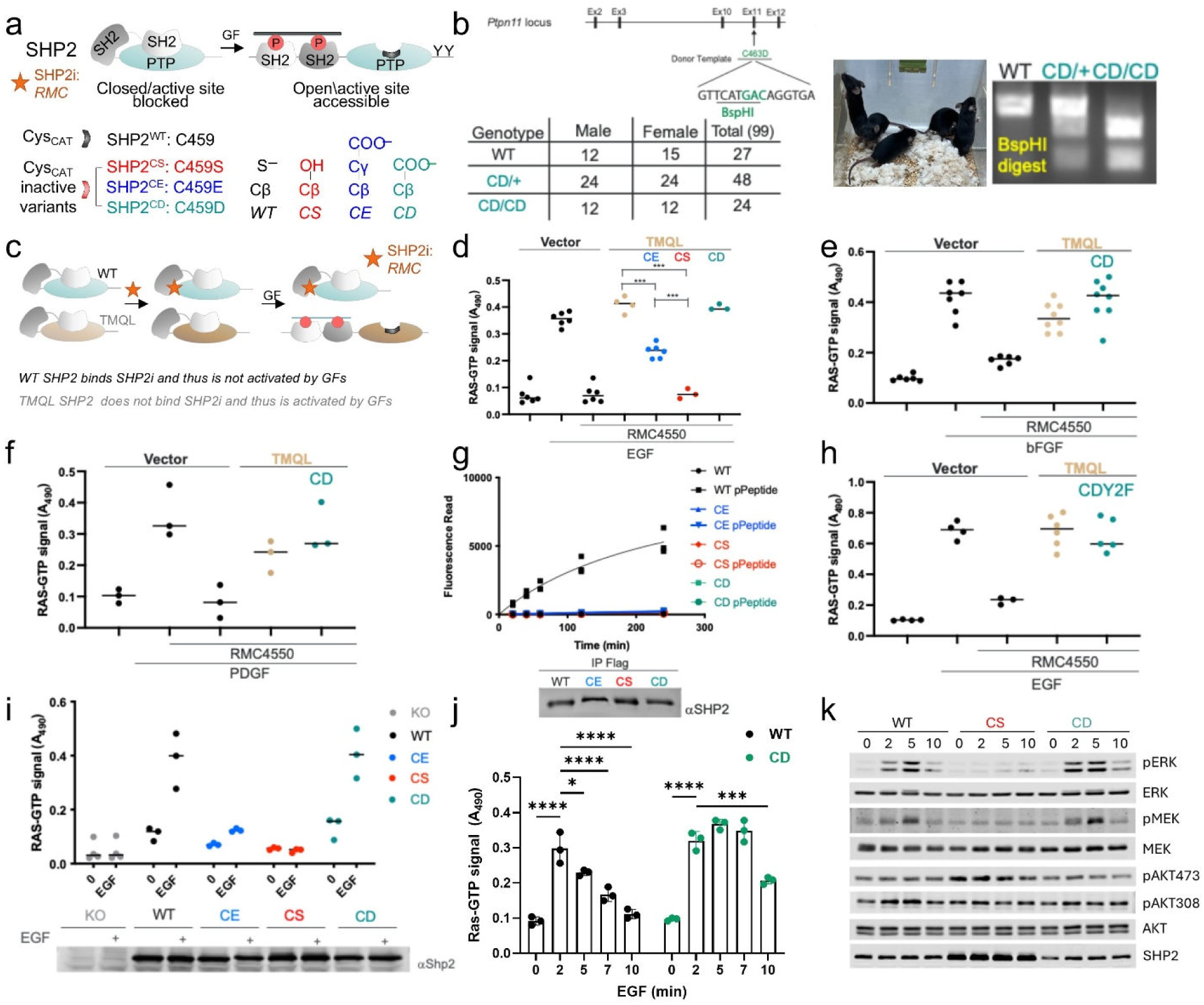
SHP2 catalytic activity is not important for RAS activation. a, SHP2 exists in either a closed, inactive (active site blocked by the SHP2 N-SH2 domain) or open, active state (N- and C-SH2 domains open to bind pY residues, making the PTP catalytic site, with the catalytic cysteine nucleophile C459, accessible). SHP2i (star), SHP2 allosteric inhibitor that stabilizes the closed state. C459 variants used in this study. **b,** Simplified targeting diagram for generating *Ptpn11^CD^* mice (upper left panel). Note that a unique restriction enzyme site was introduced to distinguish mutant and WT loci. Progeny from *Ptpn11^C463D^* matings with indicated littermates; note Mendelian inheritance (lower left panel). Picture of live animal of wild-type (WT) and *Ptpn11^CD/CD^* littermates with photograph of genotyping gel (right panel). **c,** SHP2i binds and traps SHP2^WT^ in the closed state, preventing GF-mediated activation; SHP2^TMQL^ does not bind SHP2i and thus is activated by GFs. **d-f,** RAS activation monitored by G-LISA. Lysates from 3T3 fibroblasts expressing the indicated SHP2 mutants with or without 2 min stimulation with EGF (**d**), bFGF (**e**), or PDGF(**f**), in the presence or absence of RMC4550, ***p<0.001, ANOVA. **g,** Immune complex phosphatase assay with or without activating pY peptide using DiFMUP as artificial substrate (upper panel). Similar amounts of each mutant were immunoprecipitated (lower panel). **h,** RAS activation monitored by G-LISA. Lysates from 3T3 fibroblasts expressing the indicated SHP2 mutants with or without 2 min stimulation by EGF in the presence or absence of RMC4550, as indicated. **i, j,** RAS activation in *PTPN11^-/-^* HEK293 cells reconstituted with the indicated mutants with or without EGF stimulation for 2 min (**i**) or over the indicated time course (**j**), *p<0.05, ***p<0.0005, ****p<0.0001, 2-way ANOVA. **k,** Immunoblots of lysates from *PTPN11^-/-^* HEK293 cells reconstituted with WT SHP2 or indicated mutants and stimulated with EGF for the indicated times. Representative blots from *n* = 3 biological replicates are shown.

SHP2 comprises tandem SH2 domains (N-SH2, C-SH2), a catalytic (PTP) domain, and a C-terminal tail (C-term) with two tyrosines (Y542, Y580 in humans; Y546, Y584 in mice) that recruit GRB2 upon phosphorylation (**Fig. 1a**). In the “closed” (inactive) state, the N-SH2 occludes the PTP active site preventing binding to phosphotyrosines (pYs) in RTKs or “scaffolding adapters” (e.g., GAB1-3, FRS2)^56,57^. SHP2 binding to such pY-peptides “opens” and activates SHP2 (**Fig. 1a**). This molecular switch was exploited to develop potent, selective, orally bioavailable allosteric SHP2 inhibitors (SHP2is), which bind a pocket in the closed form of SHP2 and act as “molecular glue” blocking N-SH2/loop/C-SH2 movements required for activation^1,25,32,33,58^. Hence, SHP2is effectively act as “pharmacologic protein-nulls” rather than simple catalytic inhibitors, a distinction critical to interpreting their effects on RAS/ERK signaling.

How SHP2 activates RAS has remained a mystery (**Extended Data Fig. 1a**). Multiple lines of evidence place SHP2 upstream of SOS^49,50,54,59,60^, but the exact connection is unclear. Two competing models have been proposed. The “C-terminus adaptor model” holds that SHP2 C-term pYs recruit GRB2/SOS to promote RAS exchange^61,62^. However, reconstitution studies of *Ptpn11*-mutant fibroblasts^63^ and the phenotype of mice with conditional knock-in mutations of *Ptpn11* C-terminal tyrosines show that this mechanism cannot explain SHP2 action in EGF or FGF signaling. The alternative, “substrate model”, which is supported by the effects of a small number of catalytic cysteine (Cys_CAT_) mutants (e.g. C->S, small deletions) in cell lines^37,64–67^ and in mice^68^, instead proposes that PTP activity and the dephosphorylation of (a) key substrate(s) is/are essential for RAS/ERK activation. Despite extensive efforts to identify such (a) substrate(s), no candidate can explain RTK or cytokine-evoked RAS/ERK pathway activation across signaling contexts. Furthermore, genetic support for leading candidates is lacking.

By combining cell biology, biochemistry, structural biology, advanced microscopy, and mouse and zebrafish genetics, we define how SHP2 signals to RAS. Surprisingly, we find that the SHP2 PTP domain binds directly to the SOS1 DH domain in a PTP domain-dependent but phosphatase activity-independent manner and helps recruit SOS1 to the membrane to form clusters. This interaction mediates RAS/ERK pathway activation, enables embryonic development in two vertebrate species, and facilitates transformation of hematopoietic cells to cytokine-independence. Our findings also can explain how NS and NS-ML mutants confer overlapping phenotypes despite their opposite effects on SHP2 catalysis and suggest new approaches to therapeutic targeting of SHP2.

## Results

### SHP2 catalytic activity is dispensable for RTK signaling

SHP2, like other PTP superfamily members, has an essential catalytic cysteine residue (Cys_CAT_; residue C459 in humans, C463 in mice), which acts as a nucleophile to dephosphorylate substrates via the formation of a thiophosphate intermediate^69^. If the “substrate model” is correct, homozygous Cys_CAT_ mutations and *Ptpn11* deletion should have identical phenotypic effects. Notably, Wang *et al*. recently reported that mice homozygous for the Cys_CAT_->S (CS; **Fig. 1a**) allele (*Ptpn11^CS/CS^*) are embryonic lethal (E9.5) but survive longer than *Ptpn11^-/-^* mice (E6.5-7.5)^68^. However, SHP2^CS^ is “open”^33,70,71^ and potentially “substrate trapping”^8,72^, confounding interpretation of the *Ptpn11^CS/CS^* phenotype. Therefore, we characterized the effects of two other catalytically dead variants on mouse development: Cys_CAT_->D (CD) and Cys_CAT_->E (CE) (**Fig. 1a**). Unlike SHP2^CS^, these mutants, like WT SHP2 (SHP2^WT^), adopt the closed conformation in the absence of activating pY-peptides^33,70^. As expected, the isolated SHP2^CD^ and SHP2^CE^ catalytic domains, like the SHP2^CS^ catalytic domain, showed no detectable activity against the artificial substrate 6,8-Difluoro-4-methylumbelliferyl phosphate (DiFMUP, **Extended Data Fig. 1b**). However, in contrast to the mid-gestation lethality of *Ptpn11^CS/CS^* and the peri-implantation lethality of *Ptpn11^-/-^* embryos, *Ptpn11^CE/CE^* embryos survived until E11.5-E13.5. Then they succumbed, most likely due to a markedly reduced placental labyrinth and consequent insufficiency (**Extended Data Fig. 1c**). Furthermore, *Ptpn11^CE/CE^* MEFs showed only an ∼50% decrease in EGF- and FGF-induced RAS activation compared with WT MEFs (**Extended Data Fig. 1d**). Much more surprisingly, however, *Ptpn11^CD/CD^* mice were viable and segregated in a Mendelian ratio (**Fig. 1b, Extended Data Fig. 1e**).

To compare their effects on RTK-evoked RAS activation, we generated Cys_CAT_ mutants in an SHP2i resistant background (**Fig. 1c, Extended Data Fig. 1f**; SHP2^T253M/Q257L^, hereafter, TMQL^73^). When these Cys_CAT_ mutants are expressed in the presence of an SHP2i (RMC4550, hereafter “RMC”), endogenous SHP2 is inhibited, exposing the effects of the Cys_CAT_ mutants. While SHP2^CE^ expression led to ∼50% activation, SHP2^CD^ activated RAS to similar levels as SHP2^WT^ in response to a 2-minute stimulation with EGF, FGF, or PDGF (**Figs.** **1d**-**f**). To exclude the possibility that Cys_CAT_ mutants retained PTP activity in the context of full length SHP2 (or associated with another PTP), we performed immune complex phosphatase assays (again using DiFMUP as substrate). As expected, SHP2^WT^ activity was detected in the presence of an activating pY-peptide (**Fig. 1g**); by contrast, SHP2^CD/CS/CE^ immune complexes were inactive (**Fig. 1g**). Because previous studies showed that SHP2 can auto-dephosphorylate its C-terminal pY residues, we considered the possibility that RAS activation by SHP2^CD^ might reflect increased recruitment of GRB2/SOS via the adaptor model (**Extended Data Fig. 1a**). However, there was no effect of mutating both C-terminal tyrosine residues in SHP2^CD^ (CDY2F) on growth factor-induced RAS/ERK activation (**Fig. 1h**).

Finally, we asked whether SHP2^CD^ could promote RAS/ERK activation in other cellular contexts. To this end, we reconstituted *PTPN11^-/-^* HEK cells with *PTPN11^WT^*, *PTPN11^CS^*, *PTPN11^CE^*, or *PTPN11^CD^*. As in MEFs, SHP2^CS^-expressing HEK cells had no detectable EGF-evoked RAS activation, while SHP2^CE^ enabled partial RAS activation (although to a lesser extent than in MEFs). By contrast, while initial EGF-induced RAS activation (after 2 min stimulation) was normal in SHP2^CD^-expressing cells, RAS activation was sustained (**Figs. 1i,j**). Similarly, EGF-induced MEK and ERK phosphorylation were abrogated in SHP2^CS^-expressing HEK cells but was sustained in those reconstituted with SHP2^CD^ (**Fig. 1k**). Consistent with previous reports^74,75^, pAKT-473 levels were increased in cells expressing SHP2^CS^ compared with those expressing SHP2^WT^ or SHP2^CD^. Together, these data show that SHP2 catalytic activity is dispensable for RAS/ERK activation in cells of mesenchymal (MEFs) and epithelial (HEKs) origin, as well as for development in mice.

### SHP2^CD^ is a conformational mimic of SHP2^WT^

To determine whether PTP domain-specific molecular differences in the SHP2 Cys_CAT_ mutants correlate with these unexpected outcomes, we used biomolecular NMR spectroscopy. The 2D [^1^H,^15^N] transverse relaxation optimized spectroscopy heteronuclear single quantum coherence (TROSY-HSQC, hereafter TROSY) spectra of SHP2^WT^, SHP2^CS^, and SHP2^CD^ were of high quality and suitable for analysis (**Extended Data Fig. 2a**). However, the 2D [^1^H,^15^N] TROSY spectrum of SHP2^CE^ was of low quality and careful optimization did not improve it (**Extended Data Figs. 2b,c**). Remarkably, the 2D [^1^H,^15^N] TROSY spectra of SHP2^WT^ and SHP2^CS^ showed significant chemical shift perturbations (CSPs), much beyond those expected for a single point mutation (**Figs. 2a-c**). By contrast, there were fewer and much smaller CSPs between SHP2^WT^ and SHP2^CD^ (**Figs. 2b,d,e**). Consistent with this result, the crystal structure of SHP2^CD^ is virtually identical to SHP2^WT^ (pairwise RMSDs 0.5 Å; **Extended Data Figs. 2d,e; Extended Data Table 1**). These data demonstrate that SHP2^WT^ and SHP2^CD^ adopt highly similar conformations, which are significantly different from the conformation of SHP2^CS^.

**Fig. 2.**
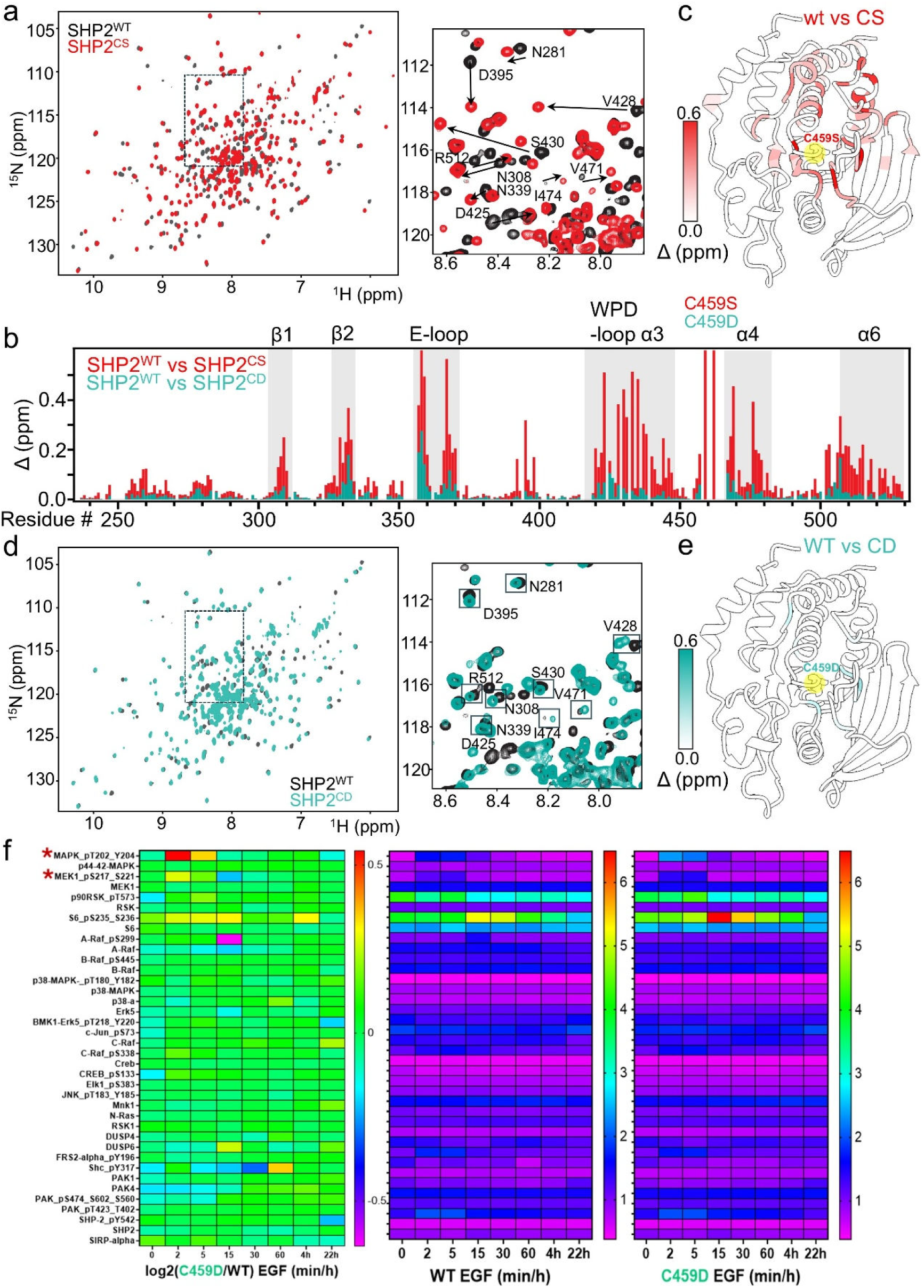
SHP2^WT^ and SHP2^CD^ are structurally and functionally highly similar. a, Overlay of 2D [^1^H,^15^N] TROSY spectra of (^2^H,^15^N)-labeled SHP2^WT^ (black) and SHP2^CS^ (red); right: zoom in of spectra with moving residues indicated. **b,** Chemical shift perturbations (CSPs) of SHP2^WT^ vs SHP2^CS^ (red) and SHP2^WT^ vs SHP2^CD^ (teal) plotted against the SHP2 protein sequence. **c,** CSPs between SHP2^WT^ and SHP2^CS^ mapped onto SHP2; C459 highlighted in yellow. **d,** Overlay of 2D [^1^H,^15^N] TROSY spectra of (^2^H,^15^N)-labeled SHP2^WT^ (black) and SHP2^CD^ (teal); right: zoom in of spectra (same as in (a)) with moving residues indicated. **e,** CSPs between SHP2^WT^ and SHP2^CD^ mapped onto SHP2; C459 highlighted in yellow. **f,** RPPA profile of MAPK pathway components from lysates of *PTPN11^-/-^* HEK293 cells reconstituted with SHP2^WT^ or SHP2^CD^ after EGF stimulation for the indicated times. Left most panel: Log_2_ ratio of normalized signal intensity values for SHP2^CD/WT^ lysates at each time point after EGF stimulation for each analyte. Middle and right panels, normalized intensity data for each analyte in SHP2^WT^ (middle) and SHP2^CD^ (right) lysates at the indicated time points after EGF stimulation. These data were used to generate the left panel. Red starts indicate the positions of p-MEK and p-ERK (aka p-MAPK).

To test if the observed similarity in SHP2^WT^ and SHP2^CD^ conformations result in globally similar effects on EGF signaling in HEK cells, we examined the effects of SHP2^WT^ and SHP2^CD^ using reverse phase protein array (RPPA) analysis. Indeed, SHP2^WT^ and SHP2^CD^ showed highly similar array results over a protracted time course following EGF addition (**Fig. 2f**), Consistent with our immunoblotting data (**Fig. 1k**), the phosphorylation of key RAS/ERK pathway components (p-MEK1, p-MAPK/ERK, p-RSK, p-S6_235, 236_) was enhanced, as was the level of the downstream pathway target DUSP6^76^, in EGF-stimulated SHP2^CD^ HEKs compared with those expressing SHP2^WT^ (**Fig. 2f**). There also was a mild overall decrease in the levels (and consequently, the phosphorylation) of many RTKs, consistent with negative feedback^77^ (**Supporting Data**). Together, these data demonstrate that SHP2^WT^ and SHP2^CD^ have similar molecular structures and evoke highly comparable signaling outcomes.

### SHP2 binds directly to SOS1

Genetic data on cell lines (e.g., shRNA, CRISPR/Cas9) show that the effects of *PTPN11* depletion are most like those of *SOS1* or *GRB2*^78,79^. Therefore, we used NMR spectroscopy to test for a potential direct interaction between the SHP2 catalytic domain (PTP; **Fig. 1a**) and GRB2 or SOS1 (**Extended Data Fig. 1a**). Comparing the 2D [^1^H,^15^N] TROSY spectrum of SHP2^CD^ with and without GRB2 showed no significant changes (**Extended Data Fig. 4a**), demonstrating that GRB2 does not bind SHP2^CD^.

We then performed similar experiments with and without various SOS1 constructs (**Fig. 3a-d**). Full length SOS1 comprises folded N-terminal regulatory (histone fold-Dbl homology-pleckstrin homology, HF-DH-PH) and catalytic (Ras exchanger motif-Cdc25, Rem-Cdc25) domains, respectively, followed by a long, disordered C-terminus that contains at least three proline rich domain binding sites for the GRB2 SH3 domains^80,81^ (**Fig. 3a**). The HF domain binds phosphatidic acid (PA), while the PH domain binds phosphatidylinositol 4,5-bisphosphate (PIP2). The Rem-Cdc25 domain contains two binding sites for RAS. Binding of RAS•GTP to the allosteric site in the Rem domain accelerates nucleotide exchange (RAS•GDP→RAS•GTP) by the Cdc25 domain (catalytic site). In cytoplasmic, inactive SOS1, the DHPH module is packed against the Rem-Cdc25 domain, obstructing RAS•GTP access to the allosteric site. Activation of SOS1 is thought to involve membrane recruitment via lipid binding to the HF and PH domains and GRB2 binding (via its SH2 domain) to activated RTKs/scaffolding adaptors/SHC with the GRB2 N-terminal SH3 domain simultaneously bound to the SOS1 C-terminal tail. However, the precise details of SOS activation remain unclear^81^. SOS1_1-1049_ lacks the disordered C-terminus but includes the HF-DH-PH and Rem-Cdc25 modules. Comparing the 2D [^1^H,^15^N] TROSY of SHP2 with and without SOS1 (increasing molar ratios of SHP2^CD^:SOS1_1-1049_) showed reductions in SHP2^CD^ resonance intensities, demonstrating that SOS1 binds directly to SHP2 (**Figs. 3b,c; Extended Data Figs. 4b,c**).

**Fig. 3.**
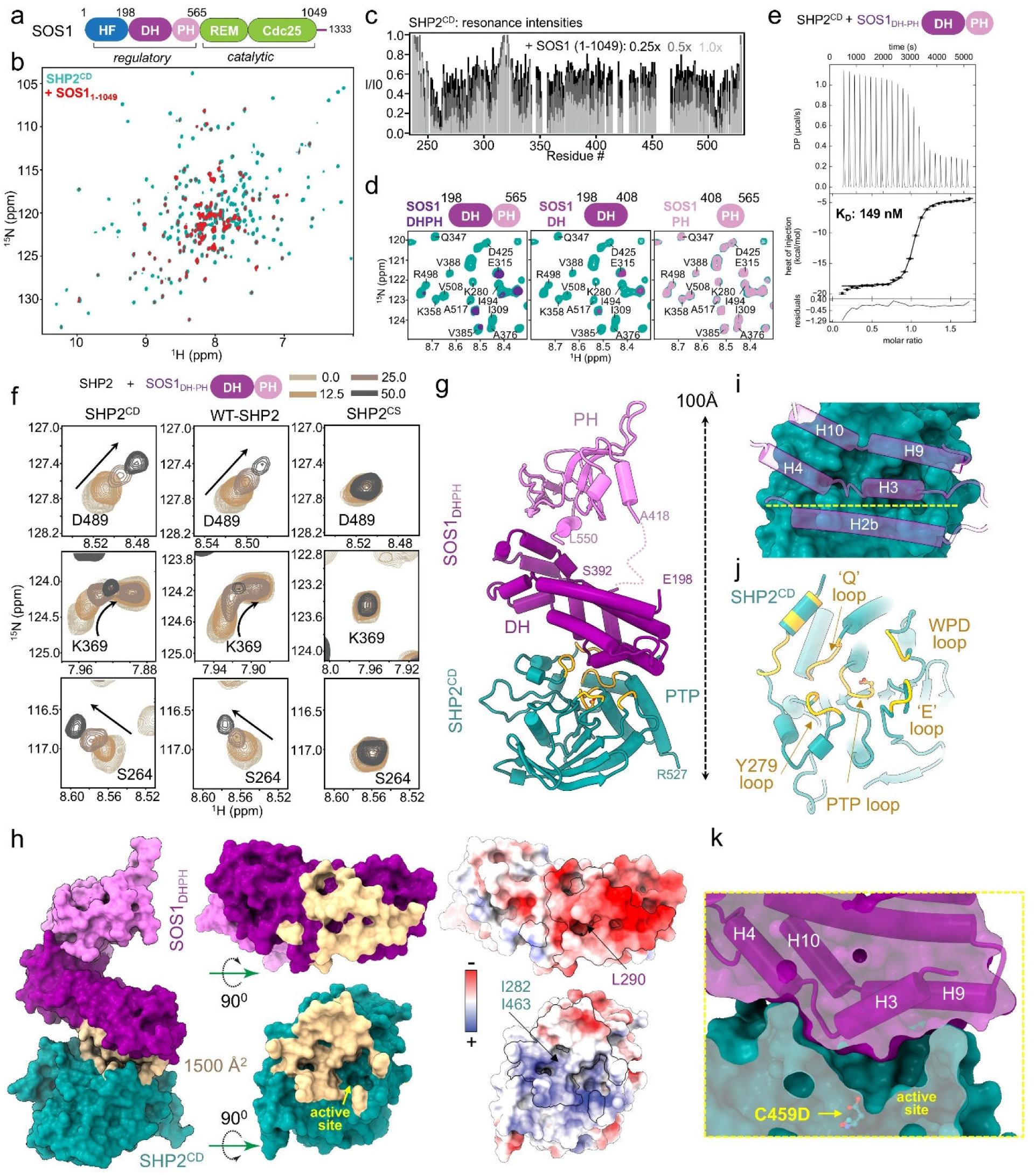
The SHP2 catalytic domain binds the SOS1 DH domain. a, SOS1 domain structure. **b,** Overlay of 2D [^1^H,^15^N] TROSY spectra of (^2^H,^15^N)-labeled SHP2^CD^ (teal) and in complex (1:1 ratio) with SOS1_1-1049_ (red). **c,** I/I_0_ intensity ratio for SOS1_1-1049_ titration (1:0.25; 1:0.5 and 1:1) plotted against the SHP2 sequence. **d,** Zoom in of 2D [^1^H,^15^N] TROSY spectra of (^2^H,^15^N)-labeled SHP2^CD^ (teal) in (1:1) complex with SOS1_DHPH_ (magenta/pink), SOS1_DH_ (magenta) and SOS1_PH_ (pink). **e,** ITC thermogram of SHP2^CD^ and SOS1_DHPH_. **f,** CSPs of SHP2 residues D489, K369 and S264 for SHP2^CD^, SHP2^WT^ and SHP2^CS^ at increasing SOS1_DHPH_ ratios. **g,** SHP2^CD^ (teal) complexed to SOS1^DHPH^(DH, magenta; PH, light pink), with N- and C-terminal residues labeled (no electron density was observed for SOS1_DHPH_ aa 393-417, and these residues were not modeled [dashed line]; orange, the SHP2 PTP loops that bind SOS1_DH_). **h,** Surface representation of the complex and individual rotated domains, with the buried residues colored beige; interface electrostatic potential shown as a surface (same orientations as those shown for buried interface) with the location of hydrophobic residues indicated. **I,** location of SOS1_DH_ helices (transparent purple) projected on SHP2 (teal). **j,** PTP domain with interface loops colored yellow and labeled. **k,** Sliced view of the SHP2^CD^:SOS1_DHPH_ interaction (slice position indicated by a yellow dashed line in *i*).

To identify the SOS1 domain(s) that mediate SHP2 binding, we repeated the binding experiments with the HF (SOS1_HF_, aa 1-198), DHPH (SOS1_DHPH_, aa 198-565), DH (SOS1_DH_, aa 198-408), PH (SOS1_PH_, aa 408-565), and Rem-Cdc25 (SOS1_cat_, aa 565-1044) domains (**Fig. 3d**, **Extended Data Fig. 4d**). Identical changes in the 2D [^1^H,^15^N] TROSY spectrum were only observed with the SOS1_DHPH_ and SOS1_DH_ domains, demonstrating that the SHP2 binds exclusively to the SOS1_DH_ domain. Furthermore, isothermal titration calorimetry (ITC) showed that SHP2^CD^ binds SOS1_DHPH_ with high affinity (K_D_ = 149 ± 66 nM; **Fig. 3e**, **Extended Data Table 2**). Consistent with the NMR and ITC data, the SOS1_DHPH:_SHP2^CD^ complex was readily purified using size exclusion chromatography (SEC; **Extended Data Fig. 5a**).

Finally, we repeated the NMR-based SOS1 interaction studies using different catalytic domain mutants of SHP2 (SHP2^WT^, SHP2^CD^, SHP2^CS^; **Fig. 3f**). Residue-specific changes in CSPs and peak intensities showed that SHP2 and SHP2^CD^ bind to SOS1_DHPH_ in a highly similar manner (**Fig. 3f, Extended Data Fig. 5b**). By contrast, the SOS1_DHPH_:SHP2^CS^ interaction was markedly attenuated, directly correlating with the effects of these mutants on RAS/ERK activation in cells (**Fig. 3f, Extended Data Fig. 5c**).

### Crystal structure of the SHP2^CD^:SOS1_DHPH_ complex

To precisely define the molecular interaction between SHP2 and SOS1, we determined the crystal structure of the SOS1_DHPH_:SHP2^CD^ complex to 2.9 Å resolution (**Fig. 3g; Extended Data Table 1**). Consistent with the NMR data (**Extended Data Figs. 5e,f**), the structure shows that SHP2^CD^ binds exclusively to the SOS1_DH_ domain on the opposite face that packs against the SOS1_PH_ domain. This elongated complex has an extensive binding interface (>1500 Å^2^ of buried surface area, **Fig. 3h**), consistent with its high affinity as determined by ITC. While most of the SHP2-interacting residues in SOS1 are from helices (**Fig. 3i**), the SOS1-interacting residues in SHP2 are largely from the loops that comprise the SHP2 catalytic pocket (**Fig. 3j**). The interacting residues are mostly charged or polar, resulting in an interface that is dominated by electrostatic interactions (**Fig. 3h**). Residues from SOS1 are mostly acidic while SHP2 residues are predominantly basic. There is a single hydrophobic interaction at the center of the interface (**Fig. 3h**). Together, these interactions anchor SOS1 helix H3 over the center of SHP2 where it binds and blocks the SHP2 active site (**Fig. 3k**). The SOS1_DHPH_:SHP2^CD^ interaction is reminiscent of the closed state complex between the N-SH2 and PTP domain of SHP2; indeed, there is substantial overlap between the SHP2 catalytic domain residues that bind the N-SH2 and SOS1_DH_ domains, respectively (**Extended Data Figs. 6a,b**). This finding predicts that only the open state of SHP2 can bind SOS1. To test this notion, we added SOS1_DHPH_ to WT (closed) or E76K (a constitutively “open” variant) SHP2_1-525_ (only lacking the C-terminal IDR tail) that includes both SH2 domains, and purified the complex by SEC. Only constitutively open SHP2 E76K, but not WT, formed a complex with SOS1_DHPH_, confirming that SHP2 must be in an open conformation to bind and recruit SOS1 (**Extended Data Fig. 6c**).

### Mutations affecting SHP2^CD^:SOS1_DHPH_ interface residues impair RAS/ERK activation but not PTP activity

The interactions that rim the outside of the SOS1-SHP2 interface are exclusively electrostatic, as are those interactions directly above the central and deep SHP2 active site **Fig. 4a-c**). The only substantial hydrophobic interaction involves SOS1_L290_, which binds a pocket defined by SHP2^I282^, SHP2^A461^, and SHP2^I463^ (**Fig. 4c**); the latter are part of the SHP2 catalytic loop (aa 457-465). To understand how these interactions contribute to RAS activation, we created single point mutations in interface residues, used retroviral transduction to express these mutants stably in *PTPN11^-/-^* HEK cells, and tested their effects on EGF-stimulated RAS/ERK activation (**Fig. 4d-g**). We chose to test SHP2 mutants so that we could also assess their catalytic activity. Indeed, as predicted, multiple mutants (F251A, Q255A, I282T, R362E, K364E, Q506K, Q506E, E508K) showed impaired or absent RAS activation (**Fig. 4d**). Except for Q255A and E508Q, which showed only a small decrease in RAS activation, these mutants also impaired ERK activation (**Fig. 4e, f**; RAS and ERK quantification compared directly in **Fig. 4g**). Remarkably, several of these mutants retained normal or even had elevated immune complex phosphatase activity (**Fig. 4h**). These findings are complementary to the effects of SHP2^CD^ on cell signaling, further highlighting that SHP2:SOS1 interaction, not SHP2 catalysis, is critical for RTK-evoked RAS activation.

**Fig. 4.**
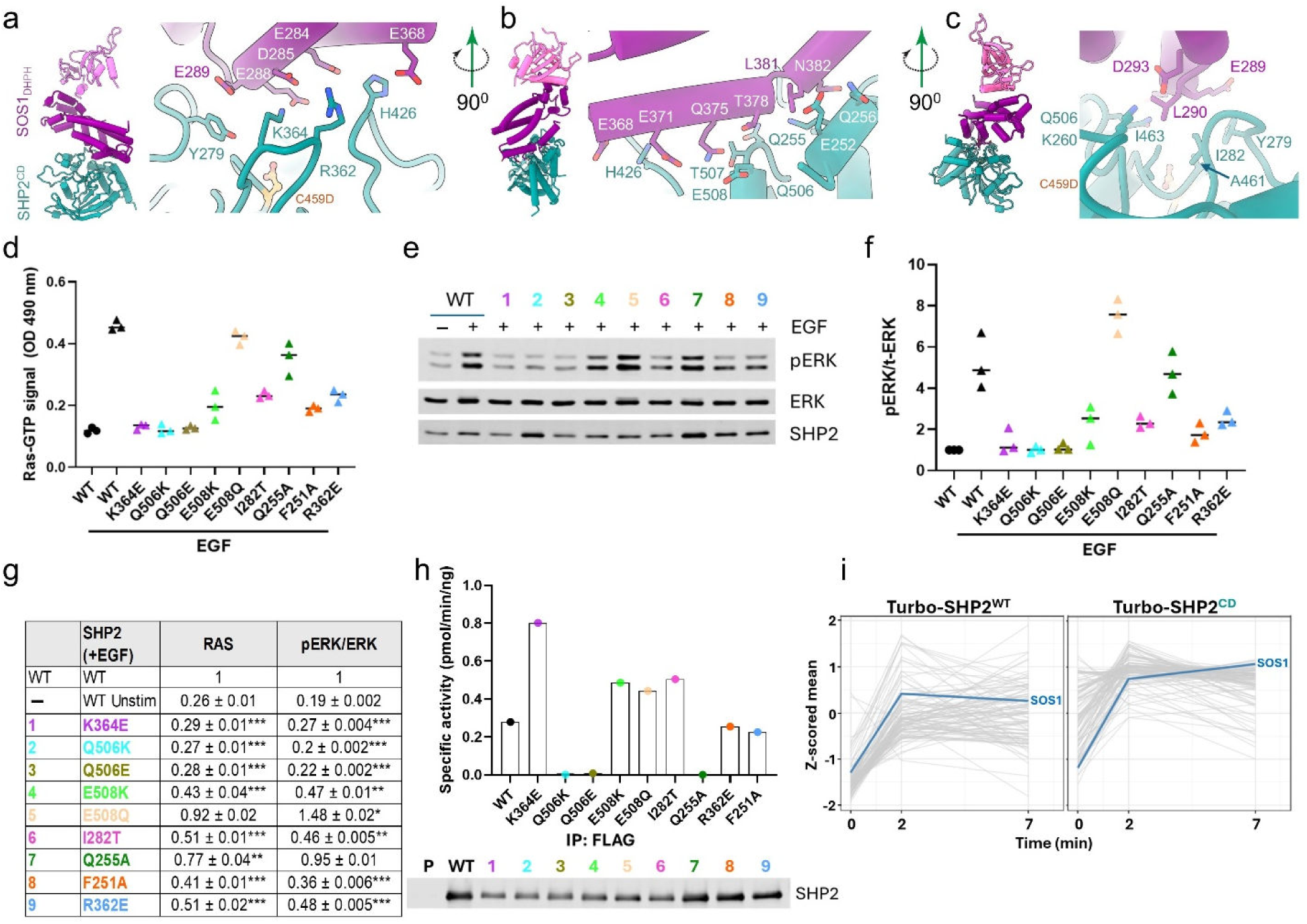
SHP2^CD^:SOS1^DHPH^ interface mutants reduce RAS activation but most are catalytically active. a-c, Zoomed in views of residues at the SHP2^CD^ (teal) and SOS1^DH^ (magenta) interface; interface residues are shown as sticks and labeled. **d,** *PTPN11^-/-^*HEK293 cells reconstituted with SHP2^WT^ or interface mutants were stimulated with EGF (25 ng/ml) for 2 min and RAS activation was quantified by G-LISA. **e, f,** Representative immunoblot (**e**) showing ERK activation in *PTPN11^-/-^* HEK293 cells with or without EGF stimulation. Blots are representative of n = 3 biologically independent experiments, which are quantified in (**f**). **g,** Statistical analysis of data from (**d**) and (**f**) illustrating RAS and ERK activation by mutant- versus SHP2^WT^-reconstituted *PTPN11^-/-^* HEK293 cells. Data are presented as mean ± s.d. for n=3 biological replicates *p<0.05, **p<0.005, ***p<0.0005, ****p<0.0001, unpaired t-test. **h,** Immune complex phosphatase performed on lysates from mutant- versus SHP2^WT^-reconstituted *PTPN11^-/-^* HEK293 cells, stimulated with EGF for 2 min. SHP2 recovery is shown below the plot. **i,** Detection of the SHP2:SOS1 interaction in cells by TurboID proximity-labeling mass spectrometry, analyzed by a linear model. Shown are Z-scored average expression values for a cluster of proteins that show increased labeling after EGF stimulation in both Turbo-SHP2^WT^ (left) and Turbo-SHP2^CD^ expressing cells (right). SOS1 is highlighted in blue. All other proteins are plotted in gray.

Our finding that mutants predicted to disrupt the SHP2/SOS1 complex impair RAS activation strongly suggested that this complex is formed in cells. Thus far, we have been unable to detect SHP2/SOS1 complexes by co-immunoprecipitation. To further assay this interaction, we generated HeLa cells expressing doxycycline (Dox)-inducible N-terminal TurboID-fusions^82^ to SHP2^WT^ and SHP2^CD^. These cells were placed in EGF- and biotin-free media for 16 hr in the presence of Dox to induce fusion protein expression. Then, biotin was added and 10 min later, the cells were stimulated with EGF for 2 or 7 min (or left unstimulated), and biotinylated proteins were recovered and analyzed by mass spectrometry (**Extended Data Fig. 6d**). Analysis of the proteomic results using a linear model revealed that labelled proteins fell into several SHP2 catalysis-dependent and catalysis-independent groups (based on whether the protein showed a similar or different pattern of behavior with SHP2^WT^ and SHP2^CD^). SOS1 was detected in a catalysis-independent group of proteins whose labeling by SHP2^WT^-Turbo and SHP2^CD^-Turbo increased following EGF stimulation with a somewhat greater increase observed for SHP2^CD^-Turbo (**Fig. 4i**). As the labeling radius of Turbo-ID is ∼10 nm^83^, these finding argue that SHP2 interacts directly with SOS1 in an EGF-dependent manner in cells.

### SHP2 catalytic domain binds SOS1 in its inactive and active conformations

The experiments above show that SHP2 in its active (open) conformation can bind to the inactive conformation of SOS1, an observation consistent with the structure of the SOS1_DHPH_:SHP2^CD^ complex, as SHP2^CD^ and the SOS1_REM_ domain bind the SOS1_DH_ on opposite faces (**Extended Data Fig. 7a**). We used NMR spectroscopy to test whether SHP2^CD^:SOS1 interaction also occurs with SOS1 in its activated, RAS-bound conformation. As noted above, this state is induced by allosteric RAS•GTP interaction with the Rem domain and subsequent conformational changes that release the Rem domain/DHPH interaction. Therefore, we tested SOS1_DHPHcat_ (SOS1 residues 198-1049), which includes the DHPH and Rem-cdc25 domains, or the membrane-independent, activation-compatible SOS1_DHPHcat:AAA_ variant E268A/M269A/D271A. The latter has a weakened PH-Rem domain interface and should render the DHPH module more accessible (“partially active”), even in the absence of RAS•GTP^84^. Upon addition of SOS1_DHPHcat_ to SHP2^CD^, robust interaction, manifested by intensity loss, was observed in the SHP2^CD^ 2D [^1^H,^15^N] TROSY spectrum (**Fig. 5a**). SOS1_DHPHcat:AAA_ addition also resulted in interaction (**Fig. 5b**). Moreover, addition of SOS1_DHPHcat:AAA_ together with HRAS^Y64A^•GppNp to induce the active state of SOS1 (RAS^Y64A^ can only bind the allosteric RAS site in SOS, not the catalytic site in the cdc25 domain^84^) also resulted in binding to SHP2^CD^ (**Fig. 5c; Extended Data Fig. 7b**). These data show that active, partially active, and inactive states of SOS1 can bind to the SHP2 catalytic domain.

**Fig. 5.**
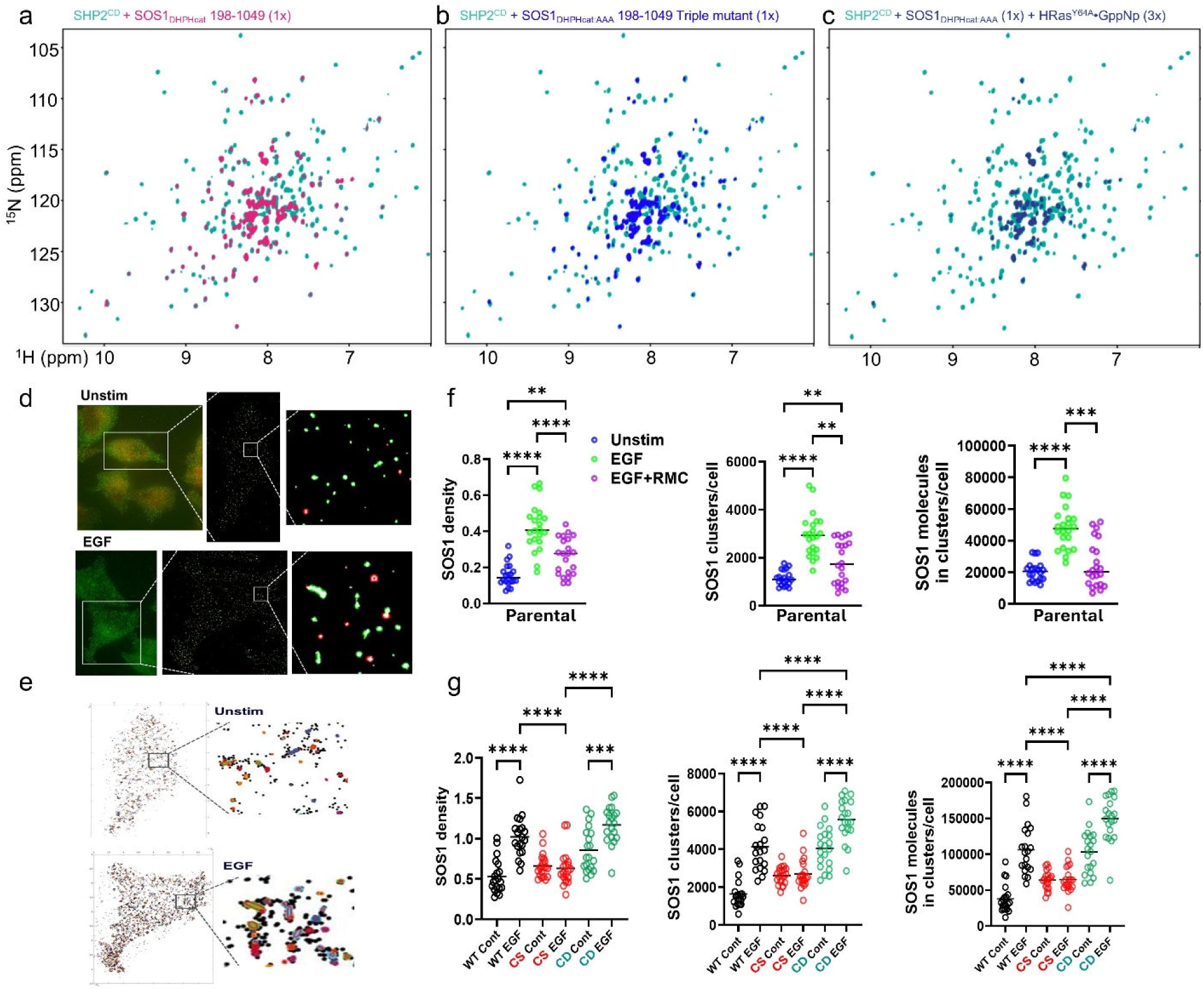
SHP2 promotes SOS1 translocation to membrane and cluster formation. a, Overlay of 2D [^1^H,^15^N] TROSY spectra of (^2^H,^15^N)-labeled SHP2^CD^ (teal) and SHP2^CD^ in complex with SOS1_DHPHcat_ (red). **b,** Overlay of 2D [^1^H,^15^N] TROSY spectra of (^2^H,^15^N)-labeled SHP2^CD^ (teal) and SHP2^CD^ in complex with SOS1_DHPHcat:AAA_ (light blue). **c,** Overlay of 2D [^1^H,^15^N] TROSY spectra of (^2^H,^15^N)-labeled SHP2^CD^ (teal) and SHP2^CD^ in complex with SOS1_DHPHcat:AAA_:HRAS^Y64A^•GppNp (dark blue). **d, e,** Representative TIRF images and reconstruction to detect localized SOS1 molecules (**d**) and clusters (**e**) in unstimulated and 2 min EGF-stimulated SOS1-Halo HeLa cells. **f, g,** Quantitative analysis of SOS1 density at the membrane (left), number of SOS1 clusters (center), and total number of SOS1 molecules in clusters per cell (right). Parental SOS1-Halo HeLa cells (**f**) or SOS1-Halo HeLa cells reconstituted with SHP2i-resistant SHP2^WT/CS/CD^, as indicated (**g**), were left unstimulated or were stimulated with EGF in the presence or absence of RMC4550 to expose mutant effects. Each data point represents analysis of an individual cell. A minimum of 20 cells per condition was analyzed in three biological replicates, **p<0.005, ***p<0.0005, ****p<0.0001, one-way ANOVA.

### SHP2 promotes SOS1 recruitment and cluster formation in response to EGF

These results raised the possibility that SHP2, via its PTP domain, might promote SOS1 recruitment to the PM. To further explore the effects of SHP2:SOS1 interaction in cells, we used super-resolution microscopy (SRM). We tagged endogenous *SOS1* in HeLa cells with Halo using CRISPR/Cas9-mediated recombination (**Extended Data Fig. 7c**). Importantly, SOS1-Halo cells showed normal RAS/ERK activation in response to EGF, indicating that tagging did not disrupt SOS1 function (**Extended Data Fig. 7d**). These cells were then starved and stimulated with EGF (2 min) in the presence or absence of RMC and imaged by Stochastic Optical Reconstruction Microscopy (STORM) in Total Internal Reflection Fluorescence Microscopy (TIRF) mode (**Fig. 5d-g**). As expected, EGF stimulation caused an increase in SOS1 recruitment to the plasma membrane (“SOS1 density”); notably, this increase was blocked by SHP2 inhibition (**Fig. 5d,f**). Interestingly, SOS1 localization was not uniform after growth factor stimulation. Instead, SOS1 formed “clusters”, defined as 3 or more SOS1 molecules within a 10 nm linking distance (**Fig. 5d-e, Extended Data Fig. 7e**). Notably, EGF stimulation increased the number of clusters per cell and the number of SOS1 molecules in clusters, and both increases were blocked by SHP2i treatment (**Fig. 5f**).

SOS1 clustering has been observed in artificial lipid bilayer systems and in T lymphocytes, where it has been associated with increased RAS exchange^85–89^. As expected, expression of inhibitor resistant SHP2 restored EGF-induced SOS1 recruitment and cluster formation in the presence of RMC (**Fig. 5g**). EGF-induced SOS1 recruitment and cluster formation were also observed in cells expressing SHP2i-resistant SHP2^CD^ (**Fig. 5g**). By contrast, inhibitor-resistant SHP2^CS^ was defective in all these functions (**Fig. 5g**). Thus, SOS1 cluster formation directly tracks RAS/ERK activation across SHP2 variants and with our structural data, suggesting that the SHP2^CD^:SOS1_DHPH_ interaction promotes the formation of multi-molecular SOS1-containing structures in response to growth factor stimulation.

### The catalysis-independent function of SHP2 is evolutionarily conserved

As noted above, *Ptpn11^CD/CD^* mice are viable and show Mendelian segregation (**Fig. 1b**). These mice are generally healthy for at least 4 months, the longest time point studied. Consistent with the mildly hypermorphic effect of SHP2^CD^ in cells, *Ptpn11^CD/CD^* mice show some mild RASopathy features (transient growth delay, altered facial proportions), which are more prominent in females (detailed phenotypic characterization of these mice will be reported elsewhere).

We next asked whether SHP2 activity is dispensable for development across evolution. Sequence analysis showed that the SHP2 and SOS1 interface residues are 100% conserved between zebrafish and humans (**Extended Data Fig. 8**). Zebrafish have two homologous *ptpn11* genes, *ptpn11a* and *ptpn11b*, of which the former is essential^90^. Although *ptpn11a* is more important for early development, *ptpn11a-/-ptpn11b-/-* fish have the most severe phenotype, characterized by absence of the swim bladder and whole body and cardiac edema. As expected, WT *zptpn11a*, but not *zptpn11a-CE,* rescued both phenotypes. Remarkably, *zptpn11a*-*CD* also restored normal early development—and to an extent comparable to WT *zptpn11a* (**Fig. 6a**).

**Fig. 6.**
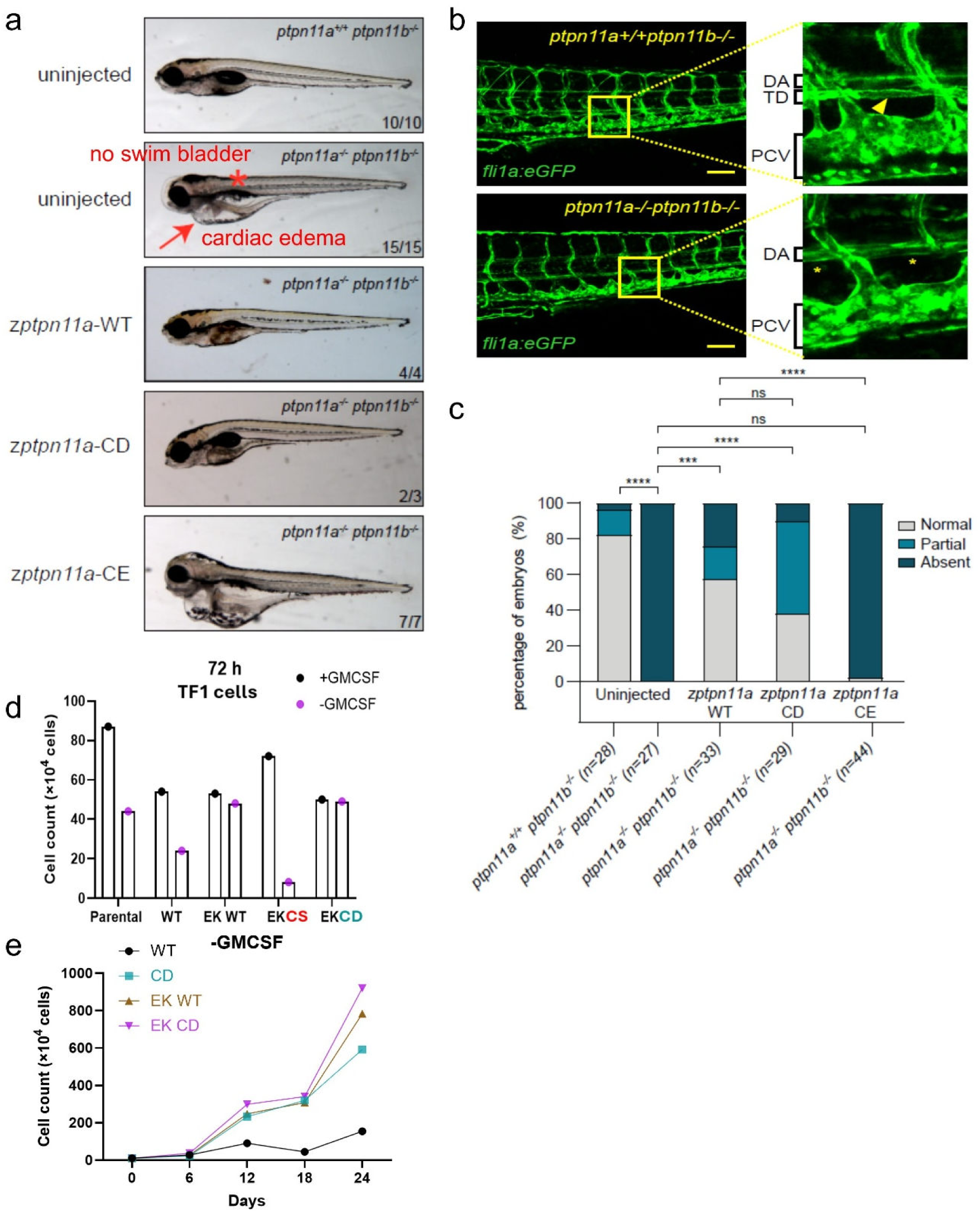
Catalysis-independent action is evolutionarily conserved and confers cytokine independence. a-c, Zebrafish embryos from an incross of *ptpn11a^+/-^ptpn11b^-/-^*fish in *Tg(fli1a:eGFP)* background were microinjected at the one-cell stage with synthetic mRNAs encoding GFP-2a-*ptpn11a* and variants. Successful injection was monitored by assessment of GFP expression at early stages. **a**, Morphology. Note absence of swim bladder and cardiac edema in *ptpn11a^-/-^ptpn11b^-/-^* embryo and rescue by *ptpn11a*-WT or *ptpn11a*-CD but not *ptpn11a*-CE. **b,** Detailed analysis of defective lymphangiogenesis phenotype. Image showing embryos at 4 dpf; note the absence of thoracic duct (TD) in *ptpn11a^-/-^ptpn11b^-/-^* embryo; dorsal aorta (DA) and posterior cardinal vein (PCV) are indicated. Scalebar is 50 µm. **c**, The number of segments with a thoracic duct was determined in 10 segments of each embryo. Embryos were then lysed, and the genotype was determined by sequencing. Embryos were scored as “normal” (TD detected in at least 7 of 10 segments), “partial” (TD detected in 1 – 6 of 10 segments), or “absent” (no TD detected in any of the segments). The results of three independent experiments are pooled with the total number of embryos (n) indicated. Statistical analysis was done using the Mann-Whitney test ***p< 0.001, ****p<0.0001 and ns = not significant. **d, e,** Proliferation assay of TF-1 cells. TF-1 cells expressing WT or mutant SHP2 were grown in the presence or absence of GMCSF. Viable cell count was determined at the indicated times by trypan blue exclusion.

Edema in *ptpn11a-/-ptpn11b-/-* embryos is caused by defective lymphangiogenesis. The full characterization of the lymphatic phenotype in zebrafish lacking functional SHP2 is described elsewhere^91^. However, a quite prominent defect is the complete absence of the thoracic duct (TD), the major lymphatic vessel in the trunk. We imaged transgenic reporter lines *Tg(fli1a:eGFP)* in which all vessels are labeled, and the TD can readily be imaged just underneath the dorsal aorta (**Fig. 6b**). We scored the presence or absence of the TD in 10 segments of the trunk per embryo and found that the TD was missing completely in *ptpn11a-/-ptpn11b-/-* embryos. Expression of WT *zptpn11a* largely rescued development of the TD, as did expression of *zptpn11-CD*, but not *zptpn11-CE* (**Fig. 6c**). The similar effects of these mutants in vertebrates separated by ∼450 million years of evolution, argues strongly that the mechanism described here is a core, ancient mechanism of RAS regulation.

### SHP2 catalytic activity is also dispensable for cytokine-independent leukemia cell growth

Leukemia-associated mutants of *PTPN11* (e.g., *PTPN11^E76K^*, *PTPN11^D61G^*) can convert normally cytokine-dependent myeloid progenitors to factor-independence. This phenomenon can be modelled *in vitro* by using TF-1 erythroblastic leukemia cells, which require GM-CSF for proliferation but not survival^92^. We engineered SHP2 CysCAT mutants into a *PTPN11^E76K^* background, transduced them into TF-1 cells, and isolated stable cell pools (**Fig. 6d**). As expected, parental TF-1 cells only proliferated in the presence of GM-CSF. All transduced lines proliferated more slowly than parental cells (in the presence of GM-CSF) because they were maintained in puromycin selection media. Upon cytokine withdrawal, SHP2^WT^-expressing cells stopped growing, whereas TF-1 cells expressing SHP2^E76K,CS^ died. As expected, SHP2^E76K^ cells proliferated equally in the presence or absence of GM-CSF but so did TF-1 cells expressing SHP2^E76K,CD^. In fact, SHP2^CD^ itself conferred cytokine-independent proliferation, consistent with its mild gain-of-function phenotype (**Fig. 6e**).

## Discussion

Over thirty years ago, Noguchi *et al.* reported that over-expression of SHP2^CS^ (“dominant negative SHP2”) impaired insulin-stimulated RAS and ERK activation in CHO-IR cells^93^. Subsequently, multiple groups, including ours, found that this mutant blocked RAS and/or ERK activation by essentially all growth factors tested (e.g., EGF, FGF, PDGF, HGF)^27,94–96^. Overexpression of SHP2^CS^ or expressing it at endogenous levels in *Ptpn11-/-* hematopoietic cells also inhibited ERK activation by several cytokines (e.g., LIF, IL3, IL2)^97–100^. Although effects on RAS activation were not assessed, SHP2 bearing a small deletion in the PTP domain impaired morphogenetic movements and FGF-induced ERK activation in *Xenopus* animal caps^65^. Mutation of a different active site residue in SHP2, the catalytic arginine (R465M) abrogated the ability of SHP2*^E76K^* to transform primary mouse bone marrow myeloid cells and impaired BCR-ABL1-induced lymphoid leukemogenesis^101,102^. Combined with findings that allosteric SHP2 inhibition blocks growth factor-evoked RAS/ERK activation and adaptive resistance, while *Ptpn11* deletion or mutation impairs vertebrate development, a consensus arose that SHP2 catalytic activity was required for the actions of SHP2 in RTK signaling, development, and oncogenesis.

Some earlier results did raise the possibility that SHP2 activity might not be absolutely required for normal or oncogenic signaling. For example, SHP2^R465M^ expression led to slightly more ERK reactivation than complete SHP2 deficiency in the setting of adaptive resistance to a KRAS inhibitor^51^. Homozygous *Ptpn11^CS/CS^* embryos were reported to live two days longer than *Ptpn11^-/-^* embryos, and MEFs from the former retained ∼50% FGF-evoked ERK activation^68^. Nevertheless, these studies were interpreted as suggesting that both SHP2 catalytic activity and its C-terminal tyrosines were necessary for signaling to the RAS/ERK pathway.

We now provide multiple lines of evidence that overturn these interpretations and explain why they were erroneous. Virtually all previous studies used SHP2^CS^ to test the role of PTP activity, and our NMR studies reveal that its active site conformation differs markedly from that of SHP2^WT^. SHP2^CD^, by contrast, is virtually indistinguishable structurally and conformationally from SHP2^WT^. Only a fidelitous structural mimic like SHP2^CD^ can isolate the effects of catalysis *per se*—and using this mutant, we find that the absence of phosphatase activity, at least on the pathways and processes tested, has surprisingly minimal consequences. Our findings also raise questions about the use of SHP2^CD^ in “substrate trapping” experiments. Although our results establish unambiguously that SHP2 catalytic activity is dispensable for most critical SHP2 functions, we do not exclude other important roles for enzyme activity. Indeed, our results suggest a likely role in RTK signal termination, and some studies implicate SHP2 in the mediating immune checkpoint inhibitor action. Future studies will be necessary to address these questions.

Instead, our results show that the SHP2 PTP domain — when not distorted (as in SHP2^CS^) — acts as a scaffold that binds the DH domain of SOS1. Multiple types of evidence support this conclusion. The SOS1_DH_:SHP2 interaction can be detected easily by NMR. ITC reveals high-affinity binding (K_D_ ≈ 150 nM) of SOS1_DHPH_ to SHP2^CD^. Proximity labeling argues that this interaction also occurs in cells. The crystal structure of the SOS1_DH_:SHP2 PTP co-complex reveals an extensive (>1,500 Å²), predominantly electrostatic, interface in which SOS1 helix H3 caps the SHP2 active site. Strikingly, multiple mutations predicted to disrupt this interface impair or abrogate growth factor-evoked RAS/ERK activation, even though some of these mutants retain WT or even elevated PTP activity. These results “square the circle”, complementing the effects of SHP2^CD^: while the latter is PTP-inactive and signaling competent/enhanced, key interface mutants are signaling impaired/dead yet PTP-active.

SOS1 is a multi-modular “coincidence detector”, which requires recruitment to the PM, lipid binding (to the HF and PH domains), disengagement of DHPH-mediated inhibition of the REM domain, and allosteric RAS•GTP binding to stimulate RAS guanine nucleotide exchange by the Cdc25 domain^81^. SHP2 PTP domain/SOS1 DH domain binding could have several potential roles in this complex process: promoting/stabilizing DH/PH domain disengagement from the REM domain to facilitate allosteric RAS binding, promoting SOS1 recruitment to/stabilization on the PM, and/or providing multivalency needed for SOS1 clustering. Our NMR data show that binding between the SHP2 PTP domain and SOS1 is comparable whether the DH/PH:REM interaction is intact or disrupted, supporting a role in recruitment. Furthermore, the extensive overlap between the DH and N-SH2 binding interfaces on the PTP domain argues that N-SH2/PTP domain interaction in “closed” SHP2 (at least in the RAS/ERK pathway) acts to prevent PTP domain/DH domain binding until SHP2 is recruited to activated, tyrosine-phosphorylated RTK or scaffolding adapters. Our SRM data provide support for a likely effect of recruitment: increasing the local concentration and multivalent interactions of SOS1 at the PM (in concert with other SOS recruiting proteins; see below), thereby promoting lipid binding to the HF and PH domains, SOS1 clustering, access to RAS•GTP, and consequently, SOS1 activation. Importantly, SOS1 clustering is known to dramatically accelerate nucleotide exchange in artificial lipid bilayers^87–89^.

In textbook models of SOS1 activation, GRB2 mediates SOS1 recruitment to RTKs and scaffolding adapters. We do not exclude such a role; indeed, in those RTK pathways where C-terminal tyrosine phosphorylation of SHP2 is partially (e.g., FGFR) or completely (PDGFR) necessary for RAS activation, we have shown that the function of these pY residues is to bind GRB2 (T. Araki *et al*., manuscript in preparation). Most likely, a critical amount of SOS1 must be recruited to localized areas on the PM to trigger SOS1 clustering and RAS activation. In RTK pathways wherein the SHP2 C-terminal tyrosines are dispensable for RAS/ERK activation, other adapters may recruit GRB2, e.g., in EGFR signaling, SHC recruits GRB2. GRB2 binds GAB1 (via its C-SH3), bringing additional SHP2 to the PM, as well as SOS1 (via its N-SH3). Together with SHP2:SOS1 binding, these interactions are likely to provide sufficient multivalency to generate the observed SOS1 clusters. Further work is required to resolve the detailed role of SHP2, SHC, and possibly other components in this process.

Our proposed “PTP scaffold” mechanism is both ancient and general. SHP2^CD^ rescues development in mouse and in zebrafish—species separated by ∼450 million years of evolution—and drives cytokine-independent proliferation of TF-1 leukemia cells. A single, conserved, catalysis-independent activity thus appears to underlie SHP2 function across normal embryonic development, RTK and cytokine signaling, and oncogenic transformation. Notably, most receptor tyrosine phosphatases (RPTPs) have an inactive “D2” domain, at least some of which retain binding capacity. Conceivably, scaffold functions arose first (in evolution) in the PTP superfamily, and catalytic function only emerged subsequently.

This revised model of SHP2 action comports with the proposed mechanism of current allosteric SHP2is, except that instead of stabilizing the closed form to prevent catalytic activation, these inhibitors prevent PTP binding to the DH domain. However, our results also identify the SHP2:SOS1 complex as an alternative target for drug design. Agents that disrupt, stabilize, or promote the degradation of this complex could quite selectively impair SHP2 action in the RAS/ERK pathway.

Finally, our findings may resolve a longstanding paradox regarding NS and NS-ML pathogenesis. NS and NS-ML mutants have opposite effects on SHP2 catalytic activity, yet the two disorders share overlapping clinical features, and both can elevate RAS/ERK output. We propose that the unifying theme of these disorders is conformational, not catalytic: both classes of mutation increase open state occupancy and should therefore enhance SOS1 binding/PM recruitment regardless of their effects on PTP activity. The mild RASopathy phenotype of *Ptpn11*^CD/CD^ is consistent with this notion. The same logic extends to the somatic *PTPN11* mutations that drive JMML, other malignancies, and temporal lobe epilepsy, and to SHP2^WT^ action in adaptive resistance. Reinterpreting SHP2 as a conformation-gated scaffold for SOS1, rather than as an enzyme whose activity is paramount in its biological and pathobiological effects, can account for the germline and somatic genetics of *PTPN11*-associated disease and suggest new ways to intervene.

## Data Availability

The NMR data generated in this study have been deposited in the BioMagResBank database under accession code BMRB 53821 (https://bmrb.io/data_library/summary/?bmrbId=53821) (SHP2), 53828 (https://bmrb.io/data_library/summary/?bmrbId=53828) (SHP2^CD^) and 53822 (https://bmrb.io/data_library/summary/?bmrbId=53822) (SHP2^CS^). The atomic coordinates and structure factors for SHP2^CD^ have been deposited in the PDB database under accession code 13KJ (10.2210/pdb13KJ/pdb). The atomic coordinates and structure factors for the SHP2^CD^:SOS1_DHPH_ complex have been deposited in the PDB database under accession code 13KI (10.2210/pdb13KI/pdb). All data (RPPA, ITC, NMR etc) generated in this study are provided in the Supplementary Information and/or in Source Data files and fully available via Figshare (https://doi.org/10.6084/m9.figshare.32661258).

## Acknowledgements

The authors thank Drs. Matthew Sale and Frank McCormick (UCSF) for helpful discussions during the early stages of this project. We thank the Perlmutter Cancer Center Rodent Genetic Engineering Laboratory resource, which is partially supported by the NYU-Perlmutter Cancer Center Support Grant (P30CA016087), for mouse targeting experiments, and Dr. Yiling Lu and Doris Siwak (MD Anderson Cancer Center) for performing RPPA. We also thank the Hubrecht Imaging Center (HIC) and animal facility of Hubrecht Institute for technical assistance. This work was supported by NIH grants R01CA49152 (B.G.N.), R01CA248896 (B.G.N. and Kwok-kin Wong), R01GM144483 and R01NS124666 (W.P.), R01GM144379 (R.P), R01MH083680 (H.S.), and U24CA270823 and U01CA271402 from the National Cancer Institute (NCI) Clinical Proteomic Tumor Analysis Consortium program (S.A.C.) and in part by the Dr. Miriam and Sheldon G. Adelson Medical Research Foundation (B.G.N., N.D.U., S.A.C., M.A.D.). Data were generated in part through the Functional Proteomics Reverse Phase Protein Array (RPPA) Core, which receives partial support from the National Cancer Institute under grant P30CA016672 to UT MD Anderson. The research reported in this publication was not directly funded through the grant P30CA016672 to UT MD Anderson and is not within the scope of such grant. This research used resources (AMX) of the National Synchrotron Light Source II, a U.S. Department of Energy (DOE) Office of Science User Facility operated for the DOE Office of Science by Brookhaven National Laboratory under Contract No. DE-SC0012704. The Center for BioMolecular Structure (CBMS) is primarily supported by the National Institutes of Health, National Institute of General Medical Sciences (NIGMS) through a Center Core P30 Grant (P30GM133893), and by the DOE Office of Biological and Environmental Research (KP1605010).

## Author Contributions

BGN, TA, WP, RP, SSK, ER, and JDH conceptualized the project and designed the experiments. TA, YA, MJG and WW performed cell biology experiments. TA performed all mouse experiments and YA and MJG all SRM studies. SSK, HLDN, and DPY expressed/purified all proteins and carried out and analyzed the NMR spectroscopy, ITC and crystallography experiments. CL, NMC, NDU, and SAC carried out and analyzed the MS proteomic studies, HS provided guidance in interpreting mouse placental phenotypes, MAD oversaw the RPPA assays, DTJW and TB performed the zebrafish studies. BGN oversaw the cell biology, mouse genetic, and in vitro biochemistry experiments. WP oversaw the NMR and ITC studies and RP supervised the crystallography studies. ER provided the equipment and oversight for SRM. JDH oversaw the zebrafish experiments. All authors contributed to writing the manuscript.

## Additional information

**Supplementary information.** The online version contains supplementary material available at XX

**Correspondence and requests for materials** should be addressed to Wolfgang Peti and Benjamin Neel.

**Reprints and permissions information** is available at http://www.nature.com/reprints.

## Methods

### Molecular cloning and mutagenesis

*PTPN11* (SHP2_237-529_, SHP2_247-529_, SHP2_237-529_C459D, SHP2_247-529_C459D, SHP2_237-529_C459S, SHP2_237-529_C459E, SHP2_1-525_, SHP2_1-526_E76K) and *SOS1* (SOS1_1-1049_, SOS1_198-1049_E268A/M269A/D271A, SOS1_198-1049_, SOS1_1-565_, SOS1_198-565_, SOS1_198-551_, SOS1_1-191_, SOS1_198-407_, SOS1_408-565_, SOS1_566-1040_) constructs for structural biology studies were cloned into either pNIC28 (MHHHHHHSSGVDLGTENLYFQS N-terminal tag) or pRP1B (MGDSKIHHHHHHENLYFQG N-terminal tag) vectors (**Extended Data Table 3**). Point mutations in *PTPN11* were introduced using QuickChange site-directed mutagenesis (Agilent). For mammalian cell expression, *PTPN11* cDNA cloned in pMSCV-IRES-GFP or pCW57.1 was used to generate mutations. Zebrafish *ptpn11a* was cloned into pCS2+ plasmids encoding eGFP followed by a peptide-2A cleavage sequence, which facilitates monitoring of expression by imaging^1^. Point mutations in *ptpn11a* were introduced by Q5 site-directed mutagenesis (New England Biolabs, E0554S).

To generate the N-terminal Myc TurboID-SHP2 fusion construct, PCR was used to amplify TurboID from V5-turboid-nes_pcdna3 (Addgene # 107169), and SHP2 (*PTPN11*) was amplified using primers containing HindIII and NotI recognition sites, a *Myc* tag and a GSGGGGSGGG linker. Primers were:

**Myc-Turbo_F:** GGTAAGCAAAGCTTGCCACCATGGAACAAAAACTCATCTCAGAAGAGGATCTCGCTAGCAAAGACAATACTGTGCCTCTG

**Turbo-linker_R:**

TCCAGACCCGCCTCCACCGGATCCCTGCAGCTTTTCGGCAGACCG

**SHP2-linker_F:**

GGTGGAGGCGGGTCTGGAGGCGGGACATCGCGGAGATGGTTTCACCC

**SHP2_R:** CTAAGCAGCGGCCGCTCATCTGAAACTTTTCTGCTGTTGCATCAGG

**Fusion_F:** GGTAAGCAAAGCTTGCCACCATG

**Fusion_R:** CTAAGCAGCGGCCGCTCA

The two fragments were combined via fusion PCR, digested with HindIII and NotI, and ligated into pcDNA5. All plasmid sequences were verified by sequencing (Quintara Biosciences or Azenta Life Sciences).

### Cell lines and virus production

Primary MEFs were prepared from E13.5 embryos and cultured in DMEM (Thermo Fisher Scientific) containing 10% FBS and 100 units/ml penicillin/streptomycin (Invitrogen), as described^2^. MEFs were starved in serum-free DMEM for 16 hrs before stimulation, then stimulated with 20 ng/ml EGF, bFGF, or PDGF (from PeproTech) before harvesting. WT MEFs were immortalized by the 3T3 method. TF-1 cells were from ATCC and were maintained in RPMI, 2 ng/ml GM-CSF (Sigma, G5035), 10% FBS, plus the above antibiotics. *PTPN11-*knockout (KO) 293 cells were generated at NCI Frederick National Laboratory for Cancer Research and were kindly provided by Drs. Matthew Sales and Frank McCormick (UCSF). They were also maintained DMEM plus 10% FBS plus antibiotics as above.

Viruses were produced by co-transfecting HEK293T cells with retroviral or lentiviral constructs and appropriate packaging vectors (Ecopack or pVSV-G with GagPol for retroviruses; pMD2.G [Addgene #12259] and psPax2 [Addgene #12260] for lentiviruses). Supernatants containing viruses were collected, passed through 0.45 μm filters, and incubated with cells for 48 hrs. After infection, transduced cells were selected by treatment with puromycin (2 µg/ml).

To assess factor dependence, parental TF-1 cells and cells expressing SHP2i-resistant *PTPN11^WT^* or various mutants were counted and seeded in the presence of RMC (3 µM) and the presence or absence of GM-CSF (2 ng/ml). The culture was replaced every 3 days, and viable cell number was counted (by trypan blue exclusion) at the indicated time points.

To generate TurboID-SHP2 knock-in HeLa cells, pcDNA5-Myc-TurboID-SHP2 constructs were inserted into FlpIn HeLa Trex cells (obtained previously from Dr. Brian Raught, Princess Margaret Cancer Center, Toronto, ON, and maintained as a Neel laboratory stock) by transfection of the pcDNA5 vector along with pOG44 (Addgene, # 209087). Transfected cells were selected in hygromycin (200 μg/ml). To generate *SOS1*-Halo knock-in HeLa cells, a targeting vector comprising a 648 bp left homology arm and a 395 bp right homology arm complementary to sequences surrounding the *SOS1* stop codon was synthesized by Twist Bioscience into the pTwist vector. Homology arms were separated by an EcoRI recognition site, which was used to insert the Halo tag in the targeting vector. The linearized targeting vector, along with a Cas9-GFP plasmid (Alt-R™ S.p. Cas9-GFP V3, IDT cat # 10008100) complexed with a *SOS1*-targeting sgRNA (CAGAGGAACTCAGGAAGAAT), were electroporated into HeLa Trex cells. Single GFP positive cells were isolated by FACS and distributed into 96-well plates. Clones screened for correct insertion by PCR, sequencing, and ultimately by immunoblotting and microscopy.

### Biochemical analyses

#### Immunoblotting

Total protein extracts from cells or tissues were prepared by homogenization in NP40 buffer (50 mM Tris-HCl pH 7.5, 150 mM NaCl, 2 mM EDTA, 1% NP40) containing a protease and phosphatase inhibitor cocktail (20 mM NaF, 1 mM Na_3_VO_4_, 10 mM β-glycerophosphate, 10 mM sodium pyrophosphate, 2 µg/ml antipain, 2 µg/ml pepstatin A, 20 µg/ml leupeptin, 20 µg/ml aprotinin, and 40 µg/ml PMSF). Homogenates were centrifuged at 16,000 x g for 10 min at 4 °C, and the supernatants were collected. Lysates were resolved by SDS-PAGE and analyzed by immunoblotting. Antibodies for immunoblots included: SH-PTP2 (B1) and ERK2 (D2) (Santa Cruz Biotechnology Inc.), and phospho-MEK1/2, MEK1/2, phospho-p44/42 MAPK, Akt1, phospho-Akt (Ser473), and pY1000 (Cell Signaling Technology). Primary antibody binding was visualized by IRDye infrared secondary antibodies using the Odyssey Infrared Imaging System (LI-COR Biosciences). Quantification of immunoblots was performed by using Odyssey V5.2.5. software. RAS activation was assessed by ELISA using a commercially available kit (Cytoskeleton; BK131). PTP activity was measured by using the full-length SHP2 assay kit (BPS Bioscience) according to the manufacturer’s instructions.

#### RPPA

SHP2-KO HEK cells reconstituted with WT SHP2 or SHP2^CD^ were serum starved for 16 hrs and lysed on ice in a buffer comprising 50 mM HEPES pH 7.4, 1% Triton-100, 150 mM NaCl, 1.5 mM MgCl_2_,1 mM EGTA,100 mM NaF, 10 mM Na pyrophosphate, 1 mM Na_3_VO_4_, 10% glycerol, and freshly added protease and phosphatase inhibitor cocktails (Roche Applied Science Cat. #s 05056489001 and 04906837001, respectively). Homogenates were centrifuged at 16,000 x g for 10 min at 4 °C, and the supernatants were collected. Protein concentration was estimated by Bradford assay and adjusted to 1.5 µg/µL using lysis buffer. The lysate was then mixed with 4 × SDS buffer (0.25 M Tris-HCl pH 6.8, 8% SDS, 40% glycerol, 10% 2-mercaptoethanol, added fresh), boiled for 5 min, and stored in -80 °C until sample submission.

Samples were processed by the RPPA Core at MD Anderson Cancer Center; details are available at https://www.mdanderson.org/research/research-resources/core-facilities/functional-proteomics-rppa-core.html. Briefly, lysates prepared as above were thawed, and five 2-fold serial dilutions were prepared in dilution lysis buffer. Serially diluted lysates were arrayed on nitrocellulose-coated slides (Grace Bio-Labs) by using a Quanterix (Aushon) 2470 Arrayer (Quanterix Corporation). A total of 5808 spots were arrayed on each slide including spots corresponding to serially diluted standard lysates, and positive and negative controls prepared from mixed cell lysates or dilution buffer, respectively. Each slide was probed with a validated primary antibody plus a biotin-conjugated secondary antibody. Antibody validation for RPPA is described on the above website. Signal detection was amplified using an Agilent GenPoint staining platform (Agilent Technologies) and visualized by DAB colorimetric reaction. Slides were scanned (Huron TissueScope, Huron Digital Pathology) and quantified using customized software (Array-Pro Analyzer, Media Cybernetics) to generate spot intensity.

Relative protein level for each sample was determined by RPPA SPACE (developed by MD Anderson Department of Bioinformatics and Computational Biology^3^). RPPA SPACE uses a logistic model to fit a single curve containing all the samples (i.e., dilution series) on a slide with signal intensity as the response variable and the dilution steps as the independent variable. The fitted curve is plotted with the signal intensities, both observed and fitted, on the y-axis and the log2 concentration of proteins on the x-axis for diagnostic purposes. The protein concentrations of each set of slides are then normalized for protein loading. Correction factor was calculated by (1) median-centering across samples of all antibody experiments; and (2) median-centering across antibodies for each sample. Results were then normalized across RPPA sets by replicates-based normalization, as described^4^.

#### Proximity labeling (Turbo-ID)/Mass Spectrometry

HeLa cells stably expressing doxycycline (Dox)-inducible *Myc*-TurboID-SHP2^WT/CD^ were serum starved for 16 h in presence of Dox (1 µg/mL). The cells were then treated with 500 µM of biotin for 10 min, during which 3 µM RMC was added for 7 min. Simultaneously, the cells were either left unstimulated or were stimulated with EGF (25 ng/mL) for either 2 or 7 mins. Following 4 washes with ice-cold PBS, cells were lysed *in situ* (on plates) on ice in RIPA buffer (50 mM Tris-HCl, pH 8, 150 mM NaCl, 5 mM EDTA, 1% NP40, 0.1% SDS) containing a protease and phosphatase inhibitor cocktail (10 mM NaF, 1 mM Na_3_VO_4_, 10 mM β-glycerophosphate, 2 µg/ml antipain, 2 µg/ml pepstatinA, 20 µg/ml leupeptin, 20 µg/ml aprotinin and 40 µg/ml PMSF). A 10% fraction of each set of beads was resolved by SDS-PAGE and silver stained to assess recovery, and the rest was sent for mass spectrometry.

Raw proteomics data were searched using Spectronaut v20.3. The protein-centric Spectronaut report was then processed using ProTIGY (https://github.com/broadinstitute/protigy-v2). Two replicates of Turbo-SHP2^CD^ at 0 minutes, one replicate of Turbo-SHP2^WT^ at 0 minutes, and one replicate of Turbo-SHP2^CD^ at 2 minutes were classified as outliers by examining the Principal Component Analysis (PCA) space and global protein intensity patterns and were excluded from downstream analyses. The remaining samples were log2-transformed, median normalized, and filtered for proteins with no more than 70% missing values across the experiment (7839/7955 proteins retained).

To identify proteins that responded to EGF stimulation in both Turbo-SHP2^WT^ and Turbo-SHP2^CD^ cells (catalysis-independent proteins), a linear model was constructed (protein expression ∼ Turbo.fusion*time) where time was treated as a factor (time = 0, 2, or 7 min). The linear model was fit to each protein using the limma package in R, and coefficients for the Turbo.fusion*time interaction and time effects were extracted. P-values for each coefficient were calculated and adjusted using the Benjamini-Hochberg correction. Catalysis-independent proteins were defined as proteins with a non-significant (adjusted p-value > 0.01 and |log_2_(FC)| < 1.25) interaction term and a significant (adjusted p-value ≤ 0.01) time term. C-means clustering was then used to group the catalysis-independent proteins by their temporal expression patterns in both Turbo-ID fusion-expressing cell populations. Proteins were grouped into 3 distinct clusters; the choice of 3 clusters was determined by minimizing the centroid distance.

#### Protein expression and purification

*PTPN11* (SHP2) and *SOS1* constructs were expressed in *Escherichia coli* BL21 (DE3) cells (Agilent). Freshly transformed cells were grown at 37°C in LB medium containing 50 µg/mL kanamycin until they reached an optical density (OD_600_) of ∼0.8. Cultures were cooled to 18°C for 1 hr, protein expression was induced by the addition of 1 mM β-D-thiogalactopyranoside (IPTG), and bacteria were allowed to grow overnight (18-20 hours, 250 rpm) at 18°C. Cells were harvested by centrifugation (6,000 *x* g, 20 min, 4°C) and stored at -80°C until purification.

For NMR experiments, expression of uniformly (^2^H,^15^N)-labeled or (^2^H,^13^C,^15^N)-labeled SHP2 (and variants) was achieved by adapting freshly transformed cells to D_2_O via multiple rounds of growth in M9 media (3 ml culture volumes) with increasing percentages of D_2_O (0%, 25%, 50%, 75%, and 100%)^5^. For protein expression, adapted cells were inoculated into 100% D_2_O M9 minimal media containing 1 g/L ^15^NH_4_Cl and/or 1 g/L ^15^NH_4_Cl and 4 g/L [^13^C-d6]-D-glucose (Cambridge Isotopes Laboratories or Isotec) as the sole nitrogen and carbon sources, respectively. The cultures were maintained at 37°C to OD_600_ of ∼1.0, then cooled to 18°C for 1 hr prior to induction by the addition of 1 mM IPTG (prepared in D_2_O). Bacteria were maintained at 18°C for 20 hrs with shaking (250 rpm), then cells were harvested by centrifugation (6,000 *x* g, 20 min, 4°C) and stored at -80°C until purification.

For all protein purifications, cell pellets were resuspended and homogenized in ice-cold lysis buffer (50 mM Tris pH 8.0, 500 mM NaCl, 5 mM Imidazole, 0.5 mM TCEP, 0.1% Triton-X-100) containing an EDTA-free protease inhibitor tablet (ThermoFisher Scientific) and lysed by high-pressure cell homogenization (Avestin Emulsiflex C3). Cell debris was pelleted by centrifugation (45,000 *x* g, 45 min, 4°C), and supernatants were filtered through 0.22 μm syringe filters (Millipore). Filtered supernatants of His_6_-tagged SHP2, SOS1, GRB2, and HRAS^Y64A^ were loaded onto a HisTrap HP column (Cytiva) pre-equilibrated with Buffer A (50 mM Tris pH 8.0, 500 mM NaCl, 5 mM Imidazole). Proteins were eluted by using a linear gradient of 0-60% Buffer B (50 mM Tris pH 8.0, 500 mM NaCl, 500 mM Imidazole). Fractions containing the protein of interest were pooled and dialyzed in TEV cleavage buffer (20 mM Tris pH 8.5, 50 mM NaCl, 1 mM TCEP) overnight at 4°C with TEV protease (prepared in-house, His_6_-tagged) to remove the N-terminal His_6_-tag. Removal of the cleaved His_6_-tag and His_6_-tagged-TEV protease was achieved by either anion exchange (SHP2 constructs) or via a ‘subtraction’ His-tag step (SOS1, GRB2, HRAS^Y64A^). Following cleavage, all SHP2 variants were loaded onto a HiTrap Q HP column (Cytiva) pre-equilibrated with Buffer C (10 mM Tris pH 8.5, 25 mM NaCl, 1 mM TCEP) and eluted using a linear gradient of 0-70% Buffer D (10 mM Tris pH 8.5, 400 mM NaCl, 1 mM TCEP). Post-cleavage SOS1/GRB2/HRAS^Y64A^ species were loaded under gravity onto Ni^2+^-NTA beads (Prometheus) pre-equilibrated with Buffer A. The flow-through and wash (buffer A) fractions, which contained the cleaved proteins were collected. Pooled protein samples were then concentrated and purified further by size exclusion chromatography (SEC; HiLoad 26/600 Superdex 200 [Cytiva]) in SEC buffer (20 mM Tris pH 7.5, 150 mM NaCl, 0.5 mM TCEP). Fractions corresponding to proteins of interest were pooled, concentrated, snap frozen in liquid nitrogen, and stored at −80°C.

#### HRAS^Y64A^ GppNp loading

For GppNp loading, a 100 µM aliquot of purified HRAS^Y64A^ protein was incubated with a 50-fold molar excess of GppNp (Sigma) in 20 mM Tris pH 8.0, 100 mM NaCl, 5 mM EDTA and 0.5 mM TCEP for 1 hr on ice. The nucleotide exchange reaction was stopped by addition of 10 mM MgCl_2_, followed by a 30 min incubation. Unbound nucleotides were removed by SEC using a Superdex 75 10/300 GL column (Cytiva; 20 mM Tris pH 8.0, 25 mM NaCl, 0.5 mM TCEP).

Nucleotide content analysis of bacterial purified as well as GppNp-loaded HRAS^Y64A^ was performed by incubating the protein at 95°C for 15 min, followed by centrifugation at 16,000 *x* g and loading of the cleared supernatant onto a MonoQ 5/50 GL (Cytiva) IEX column pre-equilibrated with 20 mM Tris pH 8.0. Anion exchange chromatography was performed by applying a linear gradient of 0-40% Buffer IEX B (20 mM Tris pH 8.0, 1000 mM NaCl, 0.5 mM TCEP) over 20 column volumes. Elution was monitored at 280/255 nm for nucleotide detection and compared against MonoQ 5/50 GL IEX chromatography runs of GDP and GppNp standards.

### NMR spectroscopy

#### Residue assignment and chemical shift perturbation measurements

All NMR data were collected on a Bruker Avance Neo 800 MHz spectrometer equipped with TCI HCN z-gradient cryoprobes at 298 K. NMR data were processed with Topspin 4.5 (Bruker) or NMRPipe^6^ and analyzed with Poky^7,8^. To assess the effects of SHP2-Cys_CAT_ mutations C459D, C459E, and C459S on PTP domain conformation, 2D [^1^H,^15^N] TROSY spectra were collected on (^2^H,^15^N)-labeled SHP2^WT/CD/CE/CS^ in NMR buffer (20 mM Na-phosphate pH 7.0, 25 mM NaCl, 1 mM TCEP and 8% D_2_O). For sequence-specific backbone resonance assignments, purified (^2^H,^13^C,^15^N)-labeled SHP2^WT/CD/CS^ were exchanged into NMR buffer and concentrated to 250, 330, or 300 µM, respectively. Backbone (H_N_, N, Cα, Cβ) chemical shift assignments were obtained by collecting triple-resonance (3D) TROSY (Transverse relaxation optimized spectroscopy) versions of HNCACB, CBCA(CO)NH, HNCA, HN(CO)CA, HNCO, HN(CA)CO experiments^9^.

The ^15^N/^1^H^N^ chemical shift perturbations (CSPs) of the WT SHP2 PTP domain vs SHP2^CD^ and SHP2^CS^ were calculated relative to WT (as reference) by comparing the 2D [^1^H,^15^N] TROSY spectra of the proteins using the equation:

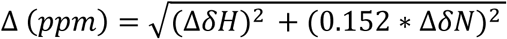

where Δ (ppm) for a given residue represents the combined chemical shift perturbation, δH represents ^1^H chemical shift difference, δN represents ^15^N chemical shift difference, and 0.152 as a ^15^N chemical shift scaling factor.

#### Detection of binding of SHP2 to GRB2 or SOS1 by NMR spectroscopy

For GRB2 binding studies, purified 50 µM (^2^H,^15^N)-labeled SHP2^CD^ was used in NMR buffer (20 mM Na-phosphate pH 7.0, 25 mM NaCl, 1 mM TCEP and 8% D_2_O) and GRB2 was added at 50 µM and 100 µM (1:1 and 1:2 molar ratio of SHP2:GRB2). For SOS1 binding studies, purified (^2^H,^15^N)-labeled SHP2^WT^, SHP2^CD^, and SHP2^CS^ PTP domains were used in NMR buffer at 50 µM and 0/12.5/25/50 µM (1:0, 1:0.25, 1:0.5 and 1:1 molar SHP2:SOS1 ratios) of SOS1 proteins: SOS1_1-1049_, SOS1_198-1049_ (SOS1_DHPHcat_), SOS1_198-1049_ E268A/M269A/D271A (SOS1_DHPHcat:AAA_), SOS1_DHPHcat:AAA_:HRAS^Y64A^•GppNp, SOS1_1-191_ (SOS1_HF_), SOS1_565-1044_ (SOS1_cat_), SOS1_408-565_ (SOS1_PH_), SOS1_198-407_ (SOS1_DH_) and SOS1_198-565_ (SOS1_DHPH_). 2D [^1^H,^15^N] TROSY spectra were collected for each sample, and resonance intensity changes in the samples to which SOS1 was added were compared to those of reference (SOS1-free) samples by calculating the intensity ratio for each residue as: I (SOS1-containing sample)/I_0_ (Reference sample). Residues with significant resonance overlaps were omitted. Additionally, SHP2 residues (D489, L283, K369, S264 and G246) manifesting detectable and progressive chemical shift changes as a function of SOS1_198-565_/SOS1_DHPH_ concentrations were identified.

#### FL SHP2:SOS1 complex binding

SHP2 and SHP2^E76K^ constructs containing the two N-terminal SH2 domains (nSH2 and cSH2), were used to assess complex formation between open (E76K) and closed (WT) states of SHP2 with SOS1_198-565_. Purified SHP2 proteins were mixed at 1:1 and 1:2 molar ratios with SOS1_198-565_ in 20 mM Tris pH 7.5, 150 mM NaCl, 0.5 mM TCEP buffer (500 µl sample volumes) and incubated on ice for 15 min. The mixtures were then analyzed by size exclusion chromatography (SEC, Superdex S75 10/300GL [Cytiva]) and column fractions were analyzed by SDS-PAGE.

### *X*- ray crystallography

#### Crystallization of SHP2_237-529_^CD^

Purified SHP2_237-529_^CD^ was concentrated to 10 mg/mL and crystallized in 0.2 M ammonium tartrate dibasic, pH 6.6, 20% PEG 3350 (w/v) in Linbro plates (Hampton Research) at a 2:1 solution:protein ratio at room temperature. Crystals were harvested, cryoprotected in 30% glycerol, and immediately flash frozen in liquid nitrogen for X-ray diffraction screening.

#### Crystallization of SHP2_247-529_^CD^:SOS1_198-551_ complex

The SHP2_247-529_^CD^:SOS1_198-551_ complex was formed by incubating the two purified proteins at 2-fold molar excess of SOS1 for 15 minutes on ice. The complex was then purified by SEC (HiLoad 26/600 Superdex 200 [Cytiva]). Fractions containing the SHP2_247-529_^CD^:SOS1_198-551_ complex were pooled, concentrated to 10 mg/ml and crystallized in 80 mM bicine, pH 9.0, 14% PEG 6000 (w/v) in Linbro plates (Hampton Research) at a 1:2 solution:protein ratio at room temperature. Crystals were harvested, cryoprotected in 30% glycerol, and immediately flash frozen in liquid nitrogen for X-ray diffraction screening.

#### Data collection, processing, and structure solution and refinement

Data processing and refinement statistics for all structures are reported in **Extended Data Table 1** with example electron density shown in **Extended Data Fig. 2e**. X-ray diffraction data for SHP2_237-529_^CD^ were collected at NSLSII (AMX beamline), and data were processed using autoProc^10^. The structure was solved by molecular replacement (MR) utilizing Phaser (v.2.8.3) in PHENIX (v.1.20.1-4487)^10^ using the structure of the SHP2 catalytic domain (PDB ID: 6ATD) as the search model. X-ray diffraction data for the SHP2:SOS1 complex were collected by using a Bruker Venture D8 system (IμS Diamond source) with a Photon III M14 detector and cold stream, and the data were processed by using PROTEUM (Bruker). The structure was solved by molecular replacement (MR) using Phaser (v.2.8.3) in PHENIX (v.1.20.1-4487) with the structures of the SOS1_DHPH_ domain (PDB ID: 1DBH) and SHP2_237-529_^CD^ (determined herein) as search models. Initial models of the complex were built using Phenix AutoBuild, followed by iterative rounds of refinement in PHENIX and manual building using Coot (v.0.9.8.7 EL)^4^.

### Isothermal Calorimetry (ITC)

ITC experiments were performed at 25°C with a Nano SV ITC microcalorimeter (TA instruments). Protein samples were dialyzed in ITC buffer (20 mM HEPES pH 7.4, 50 mM NaCl, 0.5 mM TCEP) prior to performing ITC. SHP2 was titrated into SOS1_DHPH_. Titrant (10 µL per injection) was injected into the sample cell over a period of 20 seconds with a 200-300 second interval between titrations to allow for complete equilibration and baseline recovery. Twenty-five (25) injections were delivered during each experiment, and the solution in the sample cell was stirred at 350 rpm to ensure rapid mixing. Datasets were converted using NanoAnalyze (v.3.10.0), and the converted datasets were analyzed with a one set binding site model using NITPIC (v.1.2.2)^11,12^ and SedPhat (v.12.1b)^13^. For each SedPhat evaluation the experimental parameters were set for the buffer used and the temperature, and the cell concentration was corrected to account for concentration errors^14^. Outlier data points were excluded, and the analysis was performed using the model ’A + B ↔ AB Hetero-Association’ and curve-fitting with Marquardt-Levenberg nonlinear least-squares.

### Stochastic optical reconstruction microscopy (STORM)

*SOS1*-Halo knock-in HeLa cells were used for all STORM experiments. Sample preparation was performed with fixation steps as described^15–18^. Briefly, after serum starvation for 16h, the cells were labelled with Janelia Fluor® JFX554 HaloTag® Ligand (Promega, HT 1030) for 30 min in dark. The cells were then washed three times with serum-free media and stimulated with EGF (25 ng/mL) or EGF+RMC (3 µM) for 2 min. After washing once with PBS, cells were fixed in 4% PFA+0.5% glutaraldehyde for 15 min, followed by a PBS wash and 4 washes with blocking buffer (2% glycine, 2% BSA, 0.2% gelatin, 50 mM NH4Cl). Coverslips were stored in blocking buffer overnight at 4 °C prior to imaging.

For image acquisition, the blocking buffer was replaced with <u>s</u>uper resolution (SR) imaging buffer (1 mg/mL glucose oxidase (Sigma, G2133), 0.02 mg/mL Catalase (Sigma, C3155), 10% glucose (Sigma, G8270), 100 mM mercaptoethylamine (Fisher Scientific, 100995) in PBS, pH = 8.0) to promote triplet state blinking of fluorophores and covered with a transparent seal. Samples were imaged on a custom-built optical imaging platform based on a Leica DMI 300 inverted microscope with three laser lines as described^19^. The analysis was performed via a custom MATLAB script^15–17,19^ to generate coordinates and the clustering details of individual SOS1 molecules. Python scripts were utilized to generate the localization events in individual cells, and the cell area was calculated using ImageJ. SOS1 density was calculated as total localization events/cell area. Clusters generated via the MATLAB script were defined by a linking distance of 10 nm, and the minimum number was set as 3 for each cell. At least twenty (20) cells were analyzed for each experiment, and 3 sets of independent experiments were performed. All scripts are available upon request from E. Rothenberg.

### Zebrafish experiments

All procedures involving zebrafsish were approved by the animal experiments committee of the Royal Netherlands Academy of Arts and Sciences (KNAW), Dierexperimenten commissie protocol HIdHe16596.22.01, and were conducted according to local guidelines in compliance with national and European law. Zebrafish were maintained under standard laboratory conditions^20^ and staged as described^21^. The *ptpn11a^hu3459^* and *ptpn11b^hu5920^* knockout lines were generated previously^1^. The transgenic reporter line we used in this study was *Tg(fli1a:eGFP)^y1^*^22^. Fertilized eggs were incubated at 28.5°C in E3 medium (5 mM NaCl, 0.17 mM KCL, 0.33 mM CaCl_2_, 0.33 mM MgSO_4_). 5’ capped sense mRNA was synthesized from Not1-linearized constructs using the mMessage mMachine SP6 kit (Ambion, AM1340). Embryos from an incross of *ptpn11a^+/-^ptpn11b^-/-^* zebrafish were injected at the one-cell stage with 1 nL of the mRNA solution. Scoring of the phenotypes was done by a blinded individual who assessed the morphology of embryos at 5 dpf and subsequent genotyping of individual embryos by sequencing the relevant regions of the *ptpn11a* and *ptpn11b* genes.

### Generation of Ptpn11^C463E/+^ and Ptpn11^C463D/^ mice

*Ptpn11^C463E/+^* and *Ptpn11^C463D/+^* knock-in mice were generated by *in vivo* CRISPR/Cas9 mutagenesis at the Perlmutter Cancer Center Rodent Genetic Engineering Laboratory shared resource. The sgRNA targeting exon 11 was prepared as previously described^23^. Guide DNA (GCCCTGTCGTGGTTCACTGC) was cloned into a bicistronic expression vector expressing Cas9 and sgRNA followed by addition of T7 promoter to sgRNAs template by PCR amplification. The T7-sgRNA PCR product was gel purified and used as the template for in vitro transcription using MEGAshortscript T7 kit (Life Technologies). Templates of 200 bp were centered around a C463E or C463D mutation, and transcripts were microinjected into fertilized mouse eggs from either WT or *Ptpn^LSL-D61Y/+^* female mice. High percentage chimeras were obtained, and properly targeted chimeras were crossed to C57BL/6J mice to obtain germ line transmission. For genotyping, genomic DNA was prepared from tails, then subjected to PCR and digestion with ApaLI for the C463E or BspHI for the C463D mutation. All experiments with mice were approved by the NYU Langone Institutional Animal Care and Use Committee (IACUC).

### Histology

WT and mutant placenta were obtained from timed matings. Placentas for histology were fixed in 4% paraformaldehyde at 4C overnight and embedded in paraffin. Serial sections (5µm) were prepared and stained with H&E.

### Statistics

Data are expressed as mean ± standard deviation. Statistical significance was determined using Student t test or ANOVA. Statistical analyses were performed in Prism 10 (GraphPad Software). Deviation of progeny from Mendelian frequency was assessed by χ2 test. All statistical analyses were performed with GraphPad Prism 10. For all studies, p<0.05 was considered significant.

