## Supplemental Information for "SHP2 binds directly to SOS1 to enable RAS activation"

### Extended Data for
### SHP2 binds directly to SOS1 to enable RAS activation

Toshiyuki Araki[1,#], Yashika Agrawal[1,#], Sachin S. Katti[2,#], Hiep L.D. Nguyen[2], Daniel Pushparaju Yeggoni[3], Mitchell J. Geer[1$], Wei Wei[1], Daniëlle T.J. Woutersen[4], Tieme Bijlsma[4], Cameron Genxuan Lian[5], Natalie M. Clark[5], Namrata D. Udeshi[5], Steven A. Carr[5], Heidi Stuhlmann[6], Michael A. Davies[7], Eli Rothenberg[8], Jeroen den Hertog[4,9], Rebecca Page[3], Benjamin G. Neel[1,*] & Wolfgang Peti[2,*]

[1]Perlmutter Cancer Center, NYU Grossman School of Medicine, New York, NY, USA; [2]Department of Molecular Biology and Biophysics, University of Connecticut Health Center, Farmington, CT, USA; [3]Department of Cell Biology, University of Connecticut Health Center, Farmington, CT, USA; [4]Hubrecht Institute-KNAW and University Medical Center, Utrecht, NL; [5]The Broad Institute of MIT and Harvard, Cambridge, MA, USA; [6]Department of Cell & Developmental Biology, Weill Cornell Medicine, New York, NY, USA, [7]Department of Melanoma Medical Oncology, The University of Texas MD Anderson Cancer Center, Houston, TX, USA; [8]Department of Biochemistry and Molecular Pharmacology, NYU Grossman School of Medicine, New York, NY, USA; [9]Institute Biology Leiden, Leiden University, Leiden, NL.

[$]Present address: Argonaute RNA Ltd, Science Creates - St Philips, Bristol, BS2 0XJ, UK

[#]Equal contributors to this work

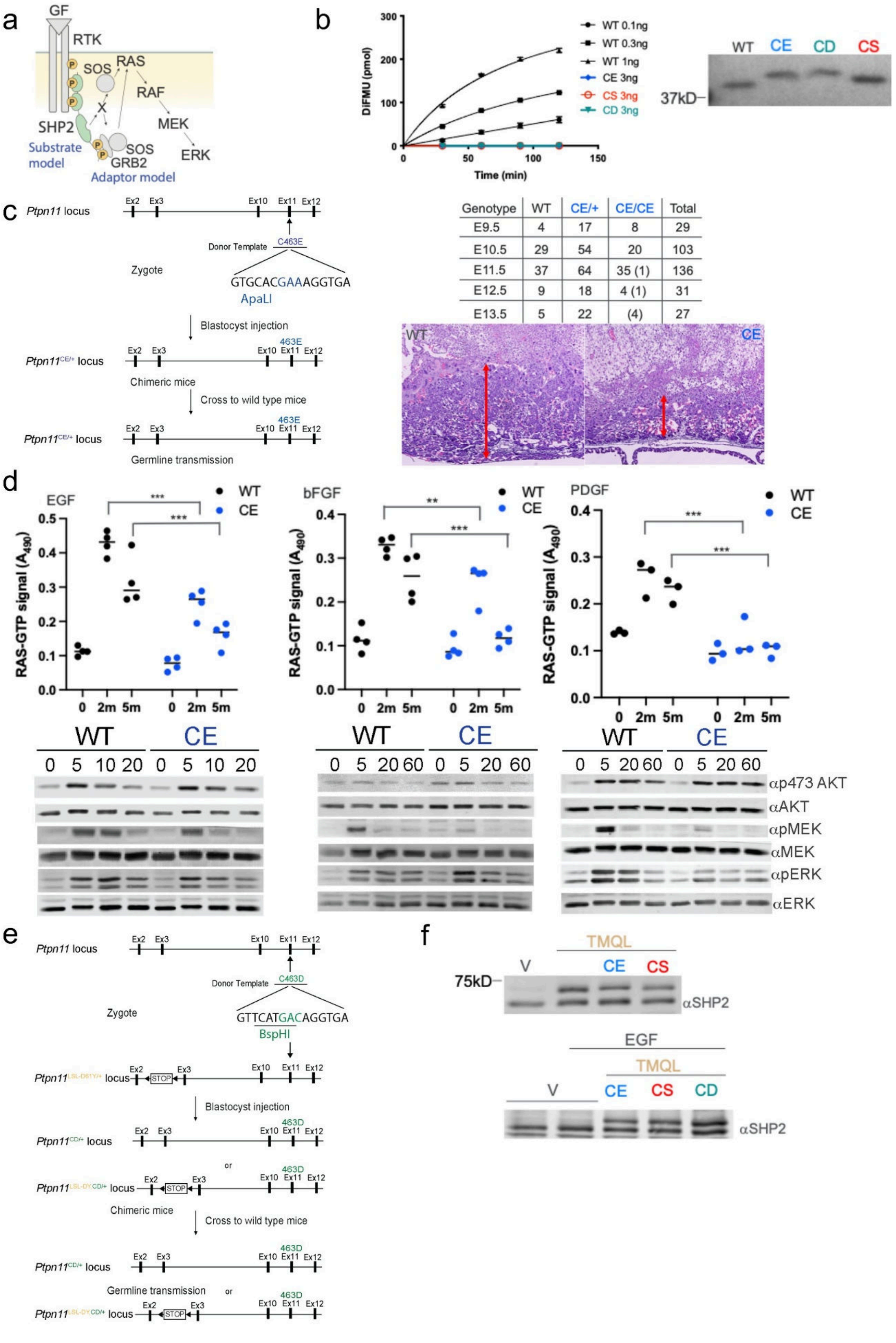

| Genotype | WT | CE/+ | CE/CE | Total |
|---|---|---|---|---|
| E9.5 | 4 | 17 | 8 | 29 |
| E10.5 | 29 | 54 | 20 | 103 |
| E11.5 | 37 | 64 | 35 (1) | 136 |
| E12.5 | 9 | 18 | 4 (1) | 31 |
| E13.5 | 5 | 22 | (4) | 27 |

**Extended Data Fig. 1│Phosphatase activity is dispensable for RAS/ERK activation by SHP2. a**, Schematic showing "adaptor" and "substrate" models for how SHP2 promotes RAS activation in response to growth factors. Tyrosine phosphorylation at C-terminus is necessary for full RAS/ERK activation by bFGF or PDGF but is dispensable for EGF. Potential substrate(s) of SHP2 is(are) indicated by "X". **b,** Phosphatase activity of isolated catalytic domains of $SHP2^{WT}$ and the indicated mutants using DiFMUP as substrate (left panel). Silver stain of SDS-PAGE gel showing proteins used for assay (right panel). **c,** Targeting strategy for *Ptpn11*$^{C463E}$ locus (left panel). Structure of *Ptpn11* locus and sequence of mutation sites on donor templates are shown. Note unique restriction enzyme sites that were introduced to detect mutations. Cas9, sgRNA targeting endogenous Cys463, and the indicated donor templates were microinjected into fertilized eggs from WT mice, followed by blastocyst injection. Chimeric offspring were crossed to WT mice to obtain germline transmission. Progeny of *Ptpn11*$^{C463E/+}$ intercrosses (right upper panel) at the indicated embryonic days; parentheses indicate dead embryos. Representative H&E-stained cross-sections of placentae from WT and *Ptpn11*$^{CE/CE}$ embryos at E11.5 (right lower panel). Note significantly smaller size of the labyrinth layer in mutant (red arrows). **d,** RAS activation (monitored by G-LISA) in WT and *Ptpn11*$^{CE/CE}$ MEFs that were left unstimulated or were stimulated with EGF (top left panel), bFGF (top middle panel) or PDGF (top right) for the indicated times. Note partial defect in activation by EGF and FGF but complete block in activation by PDGF, $^{**}p < 0.01$, $^{***}p < 0.001$, two-way ANOVA. Immunoblots of lysates that were serum-starved and stimulated with the indicated growth factors for various times and then probed with indicated antibodies (lower panels). One of three biological replicates with comparable results is shown. **e,** Details of targeting strategy for *Ptpn11*$^{C463D}$ locus. Structures of the *Ptpn11* locus and sequence of mutation sites on donor templates are shown. Note the unique restriction enzyme sites that were introduced to detect mutation. Cas9, sgRNA targeting at endogenous Cys463, and the indicated donor templates were microinjected into fertilized eggs from *Ptpn11*$^{D61Y/+}$ mice, followed by blastocyst injection. Chimeric offspring were crossed to WT to obtain germline transmission. **f,** Expression of exogenous SHP2i-resistant mutant SHP2 in 3T3 fibroblasts. Note that expression of mutants (upper bands) is similar to that of endogenous SHP2.

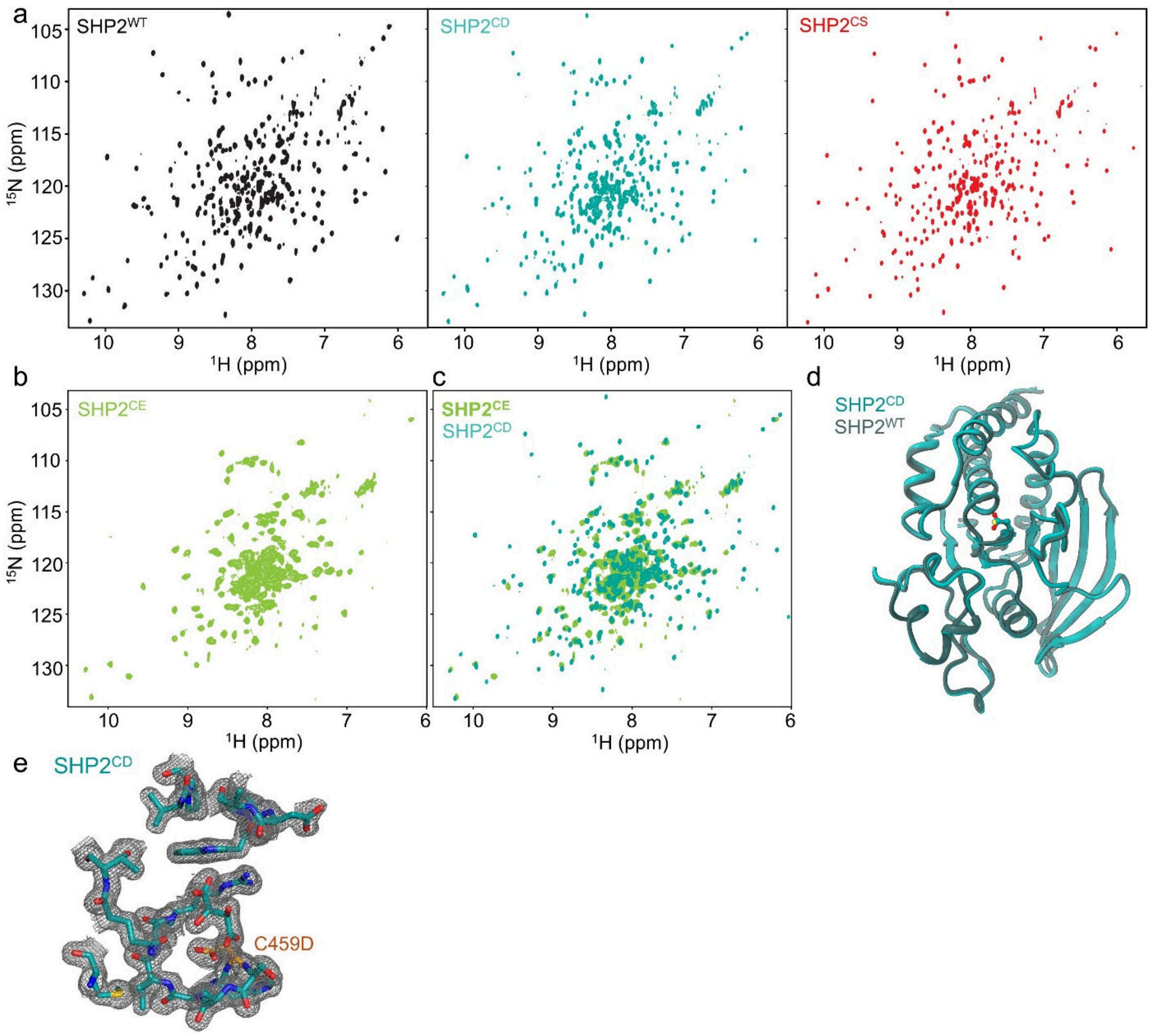

**Extended Data Fig. 2│SHP2$^{WT}$ and SHP2$^{CD}$ are structurally and functionally highly similar**. **a,** 2D [$^{1}$H,$^{15}$N] TROSY spectra of ($^{2}$H,$^{15}$N)-labeled SHP2$^{WT}$ (black), SHP2$^{CD}$ (teal) and SHP2$^{CS}$ (red). **b,** 2D [$^{1}$H,$^{15}$N] TROSY spectra of ($^{2}$H,$^{15}$N)-labeled SHP2$^{CE}$ (green). **c,** Overlay of 2D [$^{1}$H,$^{15}$N] TROSY spectra of ($^{2}$H,$^{15}$N)-labeled SHP2$^{CE}$ (green) and SHP2$^{CD}$ (teal). **d,** Superposition of crystal structure of SHP2$^{WT}$ (black) and SHP2$^{CD}$ (teal). Active site aspartic acid residue in sticks. **e,** 2$m$F$_{o}$-$D$F$_{c}$ map of SHP2$^{CD}$ crystal structure contoured at 1σ (modeled residues shown as sticks).

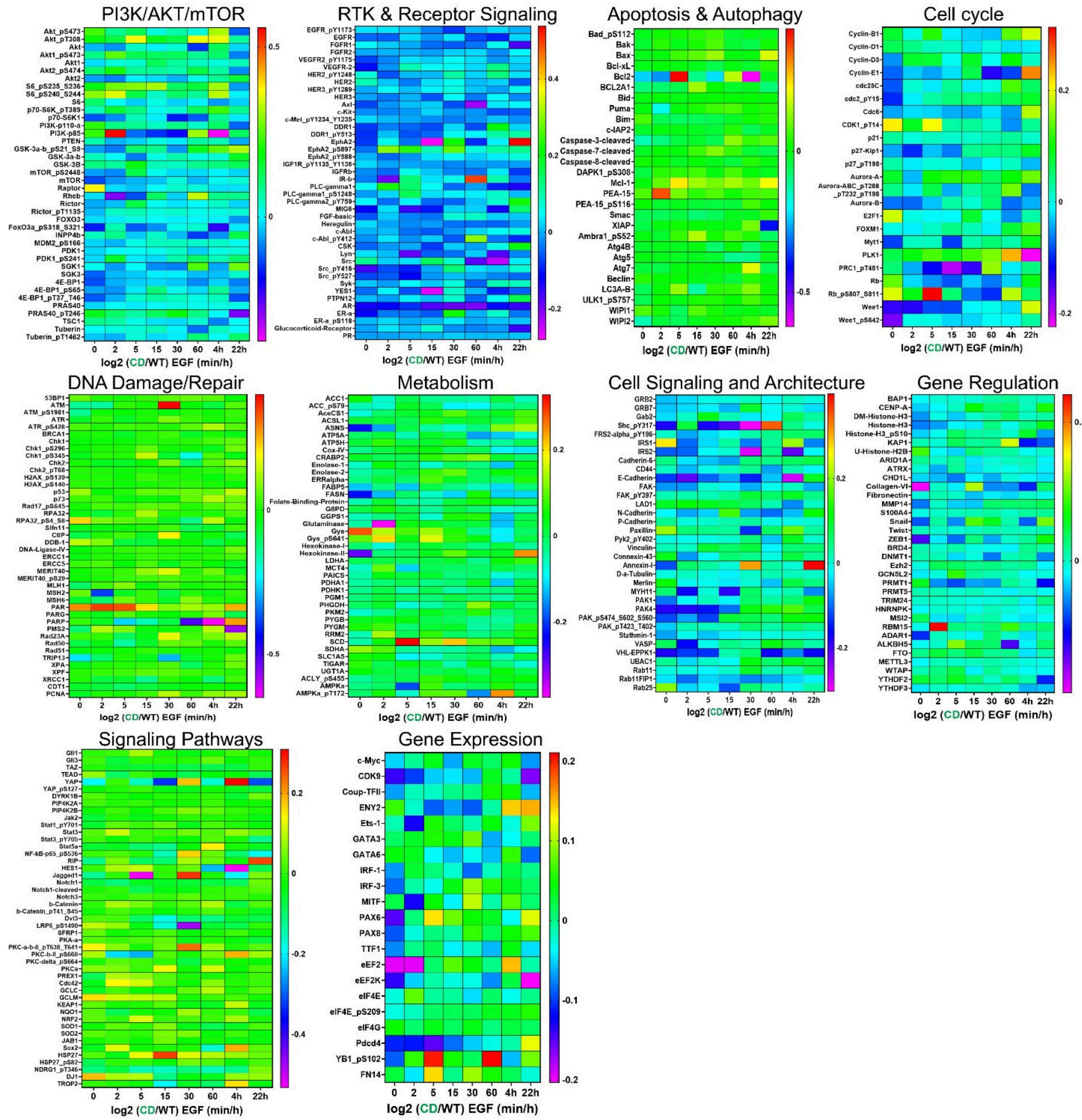

**Extended Data Fig. 3│RPPA shows similar global effects of SHP2$^{CD}$ and SHP2$^{WT}$.** RPPA profiles of different pathways, as indicated. *PTPN11*$^{-/-}$ HEK293 cells reconstituted with SHP2$^{WT}$ or SHP2$^{CD}$ were stimulated with EGF (25 ng/ml) for the indicated times and analyzed by RPPA at the MD Anderson Core Facility. Data are represented as log2 normalized intensity (SHP2$^{CD}$/SHP2$^{WT}$). Note overall global similarity of output across the time course except for downregulated "RTK and Receptor Signaling" in SHP2$^{CD}$-reconstituted cells, most likely reflecting negative feedback.

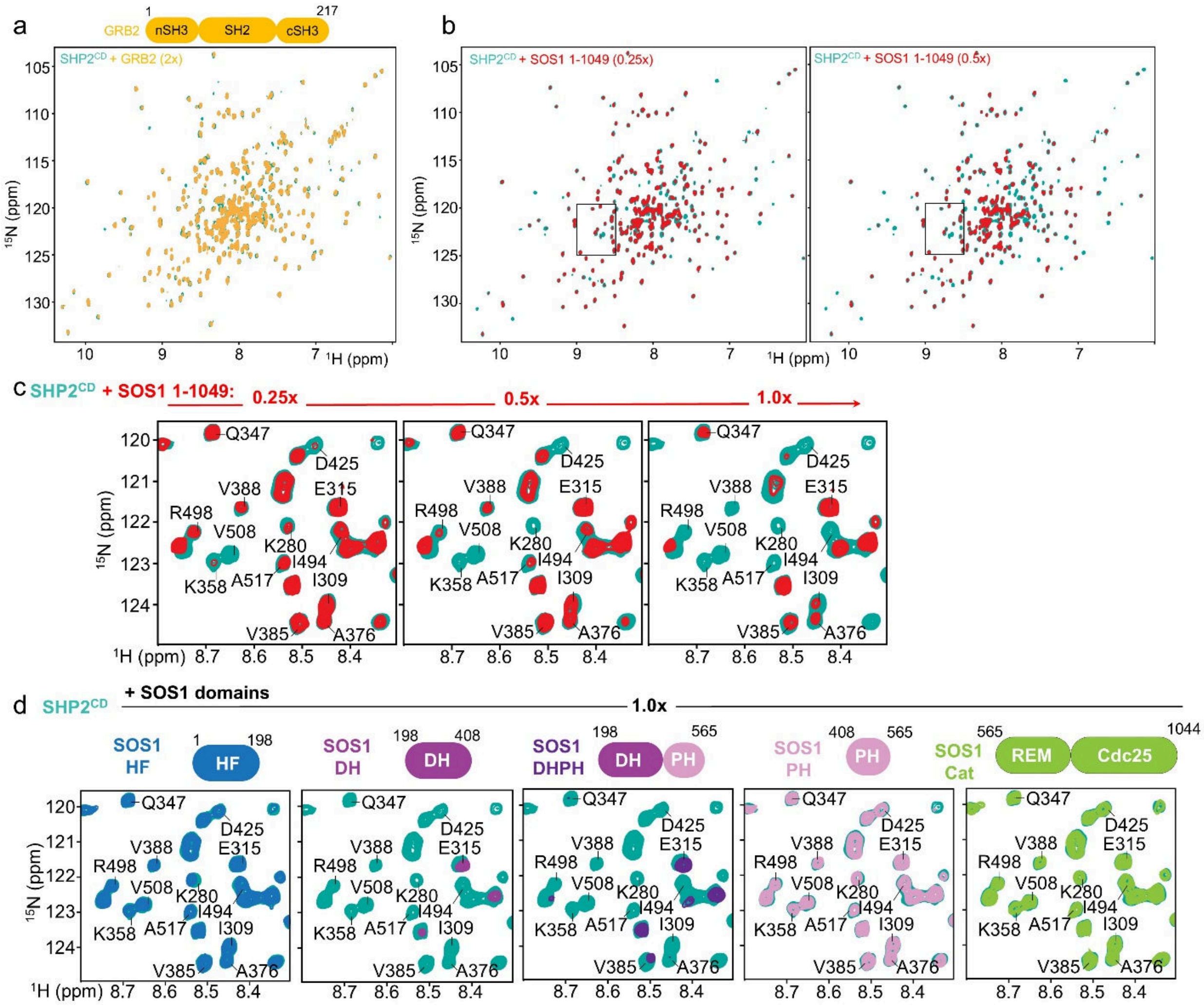

**Extended Data Fig. 4│The SHP2 catalytic domain binds the SOS1$_{DH}$ domain**. **a,** Overlay of 2D [$^1$H,$^{15}$N] TROSY spectra of ($^2$H,$^{15}$N)-labeled SHP2$^{CD}$ (teal) and in complex (1:2 ratio) with GRB2$_{1\text{-}217}$ (yellow). **b,** Overlay of 2D [$^1$H,$^{15}$N] TROSY spectra of ($^2$H,$^{15}$N)-labeled SHP2$^{CD}$ (teal) and in complex (1:0.25 and 1:0.5 ratio) with SOS1$_{1\text{-}1049}$ (red). **c,** Zoom in of 2D [$^1$H,$^{15}$N] TROSY spectra of ($^2$H,$^{15}$N)-labeled SHP2$^{CD}$ (teal) in complex (1:0.25, 1:0.5 and 1:1 ratio) with SOS1$_{1\text{-}1049}$ (red). **d,** Same zoom in as in (**c**) of 2D [$^1$H,$^{15}$N] TROSY spectra of ($^2$H,$^{15}$N)-labeled SHP2$^{CD}$ (teal) in complex (all 1:1 ratio) with SOS1$_{HF}$ (blue), SOS1$_{DH}$ (magenta), SOS1$_{DHPH}$ (magenta/pink), SOS1$_{PH}$ (pink) and SOS1$_{cat}$ (green).

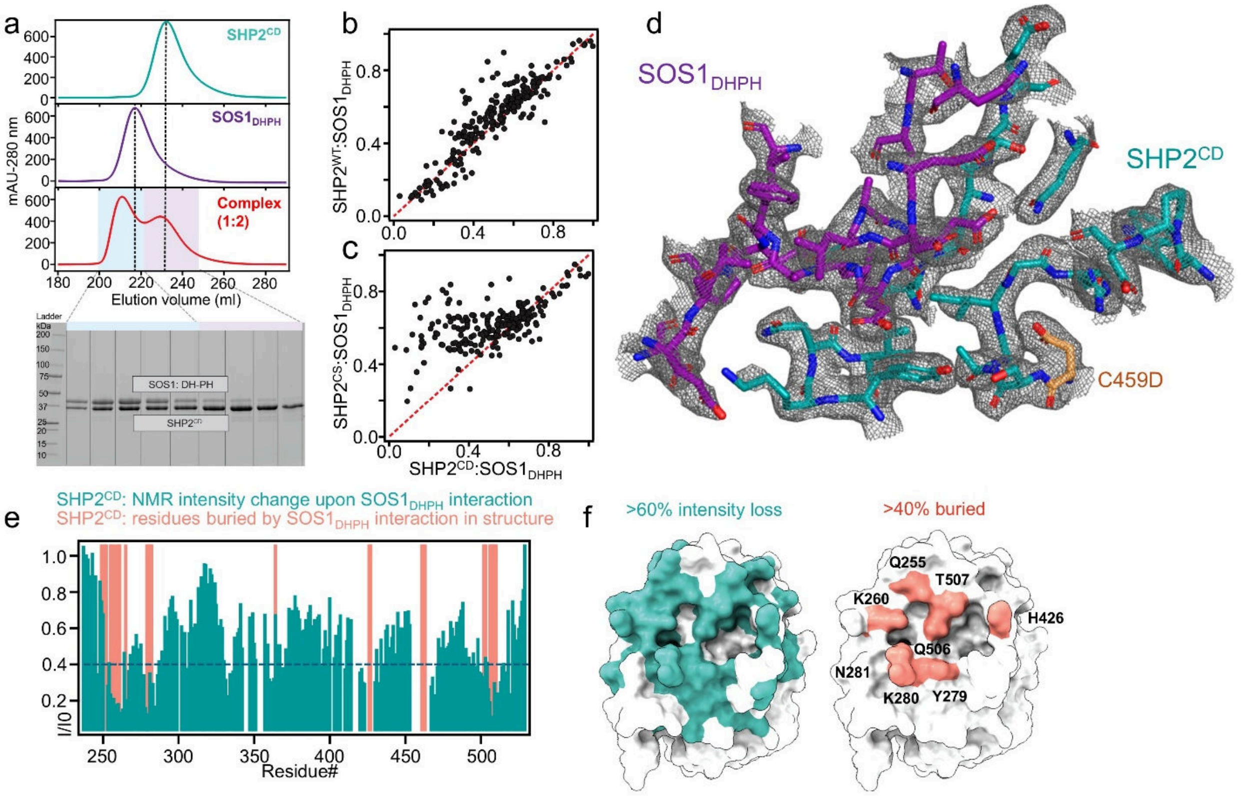

**Extended Data Fig. 5│SHP2 binds SOS1$^{DHPH}$. a,** SEC (Superdex 200 26/60) of SHP2$^{CD}$ (teal), SOS1$_{DHPH}$ (magenta) and in complex (1:2; red); below: SDS-PAGE of indicated fractions of SHP2$^{CD}$:SOS1$_{DHPH}$ complex. **b,** I/I$_0$ intensity comparison plot between SHP2$^{WT}$:SOS1$_{DHPH}$ and SHP2$^{CD}$:SOS1$_{DHPH}$. **c,** I/I$_0$ intensity comparison plot between SHP2$^{CS}$:SOS1$_{DHPH}$ and SHP2$^{CD}$:SOS1$_{DHPH}$. **d,** 2$m$F$_o$-$D$F$_c$ map of the SHP2$^{CD}$:SOS$_{DHPH}$ crystal structure contoured at 1σ (modeled residues shown as sticks). **e,** I/I$_0$ intensity comparison plot between SHP2$^{CD}$:SOS1$_{DHPH}$ vs SHP2 sequence (teal) vs newly buried SHP2 residues in SHP2$^{CD}$:SOS$_{DHPH}$ crystal structure (normalized to 1). **f,** Residues with ≥60% intensity loss (in NMR data) mapped on SHP2$^{CD}$ structure (teal) and with ≥40% BSA (in SHP2$^{CD}$:SOS1$_{DHPH}$ crystal structure) mapped on SHP2$^{CD}$ structure (key interface residues annotated).

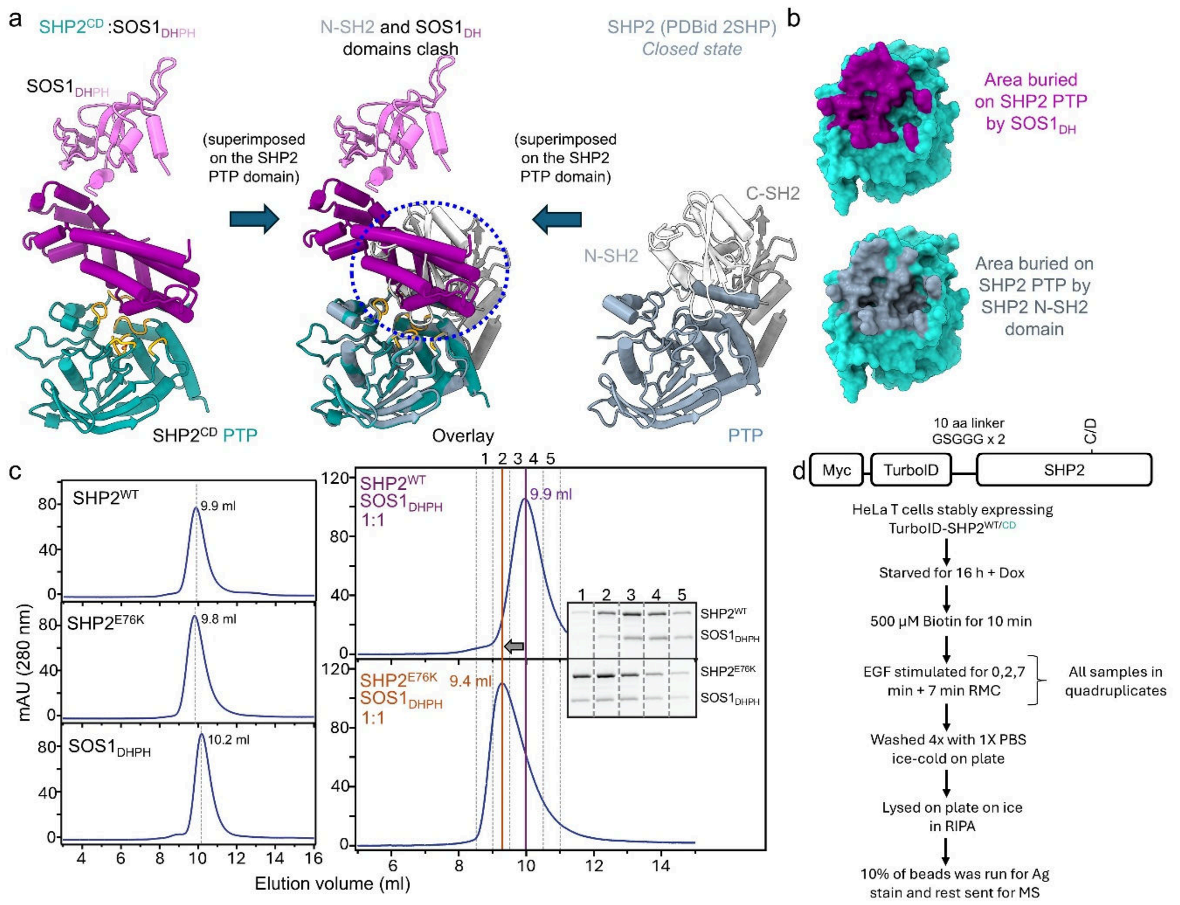

**Extended Data Fig. 6│$SOS1_{DHPH}$ binds exclusively to open SHP2. a,** Overlay (middle) of the $SHP2^{CD}$:$SOS1_{DHPH}$ complex with $SHP2^{WT}$ in the closed state (PDBid: 2SHP). **b,** Solvent accessible surface area on the SHP2 PTP domain buried by $SOS1_{DH}$ (top) or SHP2 N-SH2 (bottom). **c,** SEC (Increase 75 10/300) of full-length $SHP2^{WT}$ (top, left), $SHP2^{E76K}$ (middle, left) and $SOS1_{DHPH}$ (bottom, left) and as a 1:1 complex between $SHP2^{WT}$ and $SOS1_{DHPH}$ (top, right) or $SHP2^{E76K}$ and $SOS1_{DHPH}$ (bottom, right); inset: SDS-PAGE of indicated fractions of both complexes. **d,** Schematic of the TurboID-SHP2 genes used for proximity labeling mass spectrometry experiments (top) and scheme showing sample preparation (below).

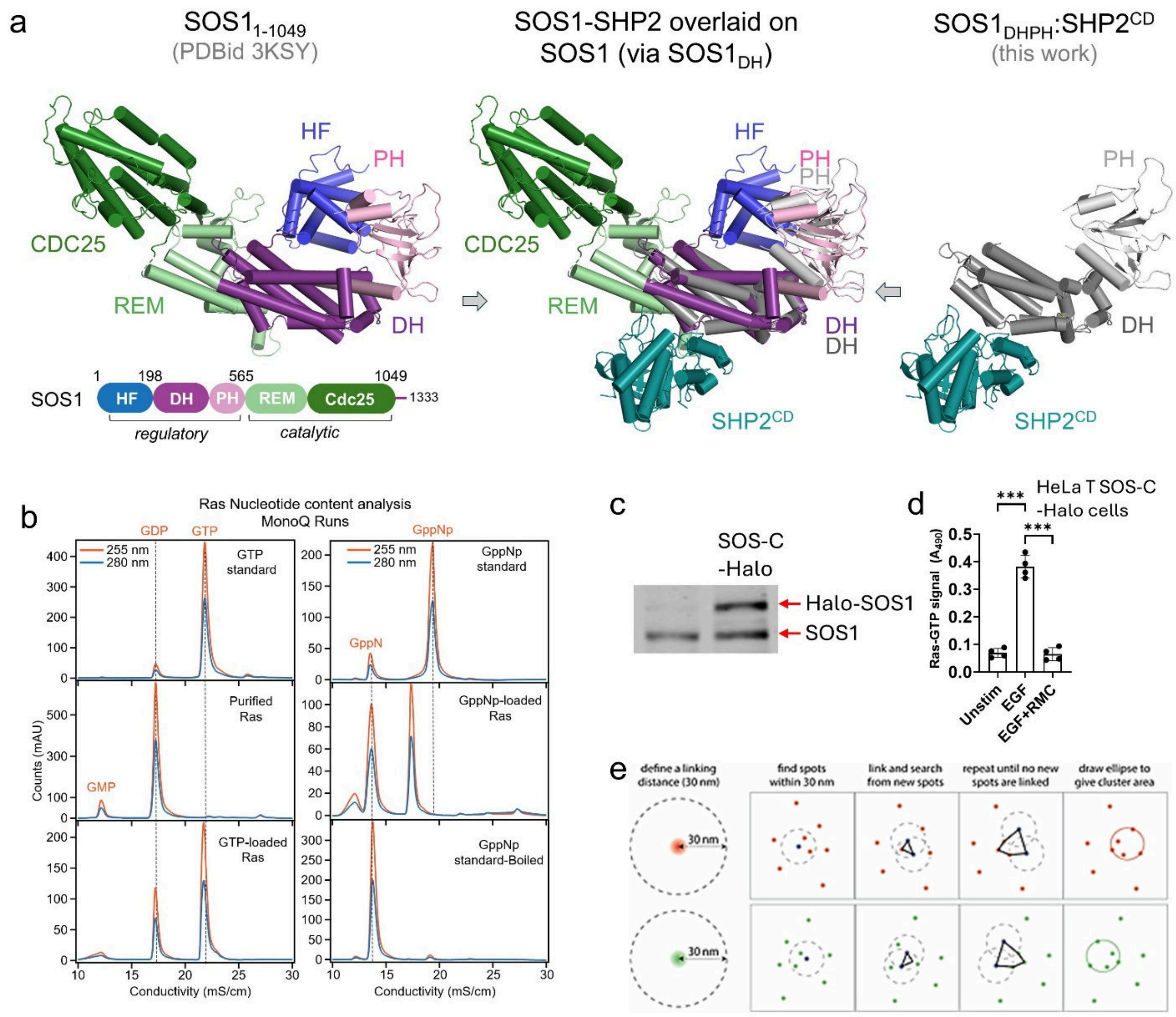

**Extended Data Fig. 7│RAS nucleotide content analysis and experimental details on super-resolution microscopy experiments. a,** Overlay of $SOS1_{1\text{-}1049}$ (left; domains colored and labeled as indicated in the cartoon; PDBid 3KSY) and the $SOS1_{DHPH}$:$SHP2^{CD}$ complex (right; $SHP2^{CD}$, teal; $SOS1_{DH}$, dark grey; $SOS1_{PH}$, light grey) via the $SOS1_{DH}$ domains from both complexes (middle), showing that $SHP2^{CD}$ and the $SOS1_{REM}$ (light green) domains bind on opposite sides of the $SOS1_{DH}$ domain. **b,** RAS nucleotide content analysis using IEX chromatography. Left: eluted standard GTP peak (top) is compared with released nucleotide peaks after boiling recombinantly purified RAS from *E. coli* (middle), and RAS•GTP (bottom). Right: eluted standard GppNp peak (top) is compared with released nucleotide peak after boiling GppNp-loaded RAS (middle). Thermal degradation of GppNp to GppN upon boiling was confirmed by loading boiled GppNp standard (bottom), with the eluted peak matching the position of GppN peak from loaded Ras (middle). **c,** SOS1 immunoblot showing successful knock-in of Halo into the endogenous *SOS1* locus of HeLa cells (right lane). Parental cells (left lane) are shown as control. **d,** RAS activation is normal in SOS1-Halo cells. SOS1-Halo HeLa cells were stimulated for 2 min with EGF in the presence or absence of RMC4550 or left unstimulated. Data shown are mean ± s.d. and are representative of 4 biological replicates, ***p<0.0005, one-way ANOVA. **e,** Schematic showing how DBSCAN algorithm defines clusters for a given linking distance.

**Sequence alignment of human SHP2 PTP domain (residues 237-529) with Zebrafish orthologs:**

```
SHP2-PTP  (Homo sapien) AETTDKVKQGFWEEFETLQQQECKLLYSRKEGQRQENKNKNRYKNILPFDHTRVVLHDGD
SHP2a-PTP (Danio rerio) AEATDKVKQGFWEEFETLQQQECKLLYSRKEGQRPENKNKNRYKNILPFDHTRVVLTDGD
SHP2b-PTP (Danio rerio) ADNSEKPKQGFWEEFEVLQQQECKLLYPRKEGQRPENKNKNRYKNILPFDTTRVQIKEAD
                        *: ::* *********.********** ****** *************** *** : :.*

SHP2-PTP  (Homo sapien) PNEPVSDYINANIIMPEFETKCNNSKPKKSYIATQGCLQNTVNDFWRMVFQENSRVIVMT
SHP2a-PTP (Danio rerio) VNEQGSDYINANLIMPDNEAKSNNSKLKRSYIATQGCLQNTISDFWRMVFQENSRVIVMT
SHP2b-PTP (Danio rerio) PDVPGSDYINANYIRSVNE-EGRHMDEGKVFIATQGCLQNTVLDFWKMVYQENTHVIVMT
                         :   ******* *    * :  .:  .: :**********: ***:**:***::*****

SHP2-PTP  (Homo sapien) TKEVERGKSKCVKYWPDEYALKEYGVMRVRNVKESAAHDYTLRELKLSKVGQGNTERTVW
SHP2a-PTP (Danio rerio) TKEVERGKSKCVKYWPDVSALKEYGAMRVRNVKETMAHDYILRELKLSKVGQGNTERTVW
SHP2b-PTP (Danio rerio) TKEMERGRNKCVRYWPDLNSTKEFGKVCVKNIEEHTAQDYIRRELEVTRLDRREPPRCIW
                        ***:***:.***:****  : **:* : *:*::*  *:**  ***:::::.: :  * :*

SHP2-PTP  (Homo sapien) QYHFRTWPDHGVPSDPGGVLDFLEEVHHKQESIMDAGPVVVHCSAGIGRTGTFIVIDILI
SHP2a-PTP (Danio rerio) QYHFRAWPDHGVPGDPGGVLDFLEEVKLKQEGITGAGPIVVHCSAGIGRTGTFIVIDILI
SHP2b-PTP (Danio rerio) HYQYLSWPDHGVPNEPGGVLSFLEQVNRTQSAIPESGPIVVHCSAGIGRTGTIIVIDILI
                        :*:: :*******.:*****.***:*: .*..*  :**:*************:*******

SHP2-PTP  (Homo sapien) DIIREKGVDCDIDVPKTIQMVRSQRSGMVQTEAQYRFIYMAVQHYIETLQRRI
SHP2a-PTP (Danio rerio) DIIREKGVDCDIDVPKTIQMVRSQRSGMVQTEAQYRFIYMAVQHYIETLQRRI
SHP2b-PTP (Danio rerio) DIINRQGLDCDIDIPKTIQRVRQQRSGMVQTEAQYKFIYMAVQQYIDTAQKRL
                        ***..:*:*****:***** **.************:*******:**:* *:*:
```

Residues interacting with SOS1 DH domain

**Sequence alignment of human SOS1 DH domain (residues 198-408) with Zebrafish ortholog:**

```
SOS1-DH (Homo sapien) EQTYYDLVKAFMAEIRQYIRELNLIIKVFREPFVSNSKLFSANDVENIFSRIVDIHELSV
SOS1-DH (Danio rerio) EQSYYELVKTFMAELRQYLRDLNLIIRVFREPFNTHTLLFSAHDVESIFSRIVDVHEVTV
                      **:**:***:****:***:*:*****:****** ::: ****:***.*******:**::*

SOS1-DH (Homo sapien) KLLGHIEDTVEMTDEGSPHPLVGSCFEDLAEELAFDPYESYARDILRPGFHDRFLSQLSK
SOS1-DH (Danio rerio) KLLGLIEDTVEMTDESSPHPLVGYCFEDLAEELAFDPYETYAQDILRSGFHEHFLSQLYK
                      **** **********.******* ***************:**:**** ***::***** *

SOS1-DH (Homo sapien) PGAALYLQSIGEGFKEAVQYVLPRLLLAPVYHCLHYFELLKQLEEKSEDQEDKECLKQAI
SOS1-DH (Danio rerio) PGAAIYLQSICEGFKEAVQYVLPRLLLTPVYHCLHYFEILKQLEEKSEDEEDKECVKQAI
                      ****:***** ****************:**********:**********:*****:****

SOS1-DH (Homo sapien) TALLNVQSGMEKICSKSLAKRRLSESACRF
SOS1-DH (Danio rerio) TALLNLQSSMERICSKSLAKRRLSESACRF
                      *****:**.**:******************
```

Residues interacting with SHP2 PTP domain

**Extended Data Fig. 8 | Human vs Zebrafish $SOS1_{DH}$ and SHP2 PTP comparison.** Sequence analysis showed that all SHP2 PTP and $SOS1_{DH}$ interface residues are 100% conserved between zebrafish and humans.

#### Extended Data Table 1: Data collection and refinement statistics

| | **$SHP2_{237\text{-}529}C459D$** | **$SHP2_{247\text{-}529}:SOS1_{198\text{-}551}$** |
|---|---|---|
| PDBID | 13kj | 13ki |
| **Data collection** | | |
| Space group | P1 | $P2_12_12_1$ |
| Cell dimensions | | |
| *a*, *b*, *c* (Å) | 39.7, 42.2, 92.0 | 56.1, 73.6, 182.8 |
| $\alpha, \beta, \gamma$ (°) | 94.6, 92.6, 108.5 | 90, 90, 90 |
| Resolution (Å) | 39.81 – 1.63 (1.71 – 1.63) | 21.82 – 2.88 (3.05 – 2.88) |
| $R_{merge,}$ | 0.09 (1.26) | 0.23 (1.35) |
| $R_{meas}$ | 0.10 (1.48) | 0.24 (1.50) |
| $I/\sigma(I)$ | 7.2 (1.1) | 7.9 (1.2) |
| $CC_{1/2}$ | 0.99 (0.46) | 0.99 (0.48) |
| Completeness (%) | 97.5 (96.1) | 99.5 (98.8) |
| Redundancy | 3.7 (3.7) | 7.8 (5.6) |
| **Refinement** | | |
| Resolution (Å) | 37.91 – 1.63 (1.67 – 1.63) | 21.82 - 2.88 (2.97-2.88) |
| No. reflections | 62357 (1799) | 16150 (1066) |
| $R_{work}$ / $R_{free}$ | 0.21 (0.24) | 0.24 (0.28) |
| No. atoms | | |
| Protein | 4483 | 4639 |
| Ligand/ion | 20 | -- |
| Water | 336 | 11 |
| *B* factors | | |
| Protein | 26.9 | 56.0 |
| Ligand/ion | 36.5 | -- |
| Water | 31.7 | 28.8 |
| R.M.S. deviations | | |
| Bond lengths (Å) | 0.005 | 0.002 |
| Bond angles (°) | 0.700 | 0.466 |

Values in parentheses are for highest-resolution shell

**Extended Data Table 2: ITC $SOS1_{DHPH}$:SHP2**

| Cell | Titrant | $K_d$ (nM) | ΔH (kcal/mol) | TΔS (kcal/mol) | ΔG (kcal/mol) | n |
|---|---|---|---|---|---|---|
| $SOS1_{DHPH}$ | $SHP2^{CD}$ | 149 ± 66 | -16.2 ± 2.3 | -6.4 ± 1.7 | -9.4 ± 0.1 | 6 |

*All experiments were performed at 25 °C. Values are reported as mean ± standard deviation for experiments (n≥3). $SOS1_{DHPH}$ (aa 198-565); SHP2 (aa 237-526)

**Extended Data Table 3: Constructs used in this study**

| pRP1B vector constructs | pNIC28 vector constructs | pMSCV-IRES-GFP vector constructs | pCW57.1 vector constructs |
|---|---|---|---|
| $SOS1_{1-1049}$ (HF-DH-PH-cat) | WT $SHP2_{237-529}$ | $SHP2_{1-593}$ T253M/Q257L | $SHP2_{1-593}$ T253M/Q257L |
| $SOS1_{198-1049}$ (DH-PH-cat) | WT $SHP2_{247-529}$ | $SHP2_{1-593}$ TMQL/C459E | $SHP2_{1-593}$ TMQL/C459S |
| $SOS1_{198-1049}$ E268A/M269A/D271A | $SHP2_{237-529}$ C459S | $SHP2_{1-593}$ TMQL/C459S | $SHP2_{1-593}$ TMQL/C459D |
| $SOS1_{1-565}$ (HF-DH-PH) | $SHP2_{237-529}$ C459D | $SHP2_{1-593}$ TMQL/C459D | TurboID-$SHP2_{1-593}$ |
| $SOS1_{198-565}$ (DH-PH) | $SHP2_{247-529}$ C459D | $SHP2_{1-593}$ TMQL/C459D/ Y542F/Y580F | TurboID-$SHP2_{1-593}$ C459D |
| $SOS1_{198-551}$ (DH-PH) | $SHP2_{237-529}$ C459E | $SHP2_{1-593}$ TMQL/E76K | |
| $SOS1_{1-191}$ (HF) | | $SHP2_{1-593}$ TMQL/E76K/ C459S | |
| $SOS1_{198-407}$ (DH) | | $SHP2_{1-593}$ TMQL/E76K/ C459D | |
| $SOS1_{408-565}$ (PH) | | $SHP2_{1-593}$ | |
| $SOS1_{566-1040}$ (cat) | | $SHP2_{1-593}$ C459E | |
| WT $SHP2_{1-525}$ | | $SHP2_{1-593}$ C459S | |
| $SHP2_{1-526}$ E76K | | $SHP2_{1-593}$ C459D | |
| $GRB2_{1-217}$ | | $SHP2_{1-593}$ K364E | |
| H-$RAS_{1-166}$ Y64A | | $SHP2_{1-593}$ Q506K | |
| | | $SHP2_{1-593}$ Q506E | |
| | | $SHP2_{1-593}$ E508K | |
| | | $SHP2_{1-593}$ E508Q | |
| | | $SHP2_{1-593}$ I282T | |
| | | $SHP2_{1-593}$ Q255A | |
| | | $SHP2_{1-593}$ F251A | |
| | | $SHP2_{1-593}$ R362E | |
